# Female rat sexual behavior is unaffected by perinatal fluoxetine exposure

**DOI:** 10.1101/2020.05.29.122945

**Authors:** Jan Hegstad, Patty T. Huijgens, Danielle J. Houwing, Jocelien D.A. Olivier, Roy Heijkoop, Eelke M.S. Snoeren

## Abstract

Serotonin plays an important role in adult female sexual behavior, however little is known about the influence of serotonin during early development on sexual functioning in adulthood. During early development, serotonin acts as neurotrophic factor, while it functions as a modulatory neurotransmitter in adulthood. The occurrence of serotonin release, could thus have different effects on behavioral outcomes, depending on the developmental period in which serotonin is released. Because serotonin is involved in the development of the HPG axis which is required for puberty establishment, serotonin could also alter expression patterns of for instance the estrogen receptor α (ERα).

The aim of our study was to investigate the effects of increased serotonin levels during early development on adult female rat sexual behavior during the full behavioral estrus in a seminatural environment. To do so, rats were perinatally exposed with the selective serotonin reuptake inhibitor (SSRI) fluoxetine (10 mg/kg FLX) and sexual performance was tested during adulthood. All facets of female sexual behavior between the first and last lordosis (behavioral estrus), and within each copulation bout of the behavioral estrus were analyzed. Besides the length and onset of the behavioral estrus and the sexual behaviors patterns, other social and conflict behavior were also investigated. In addition, we studied the effects of perinatal FLX exposure on ERα expression patterns in the medial preoptic nucleus, ventromedial nucleus of the hypothalamus, medial amygdala, bed nucleus of the stria terminalis, and the dorsal raphé nucleus.

The results showed that perinatal fluoxetine exposure has no effect on adult female sexual behavior. The behavioral estrus of FLX-females had the same length and pattern as CTR-females. In addition, FLX- and CTR-females showed the same amount of paracopulatory behavior and lordosis, both during the full behavioral estrus and the “most active bout”. Furthermore, no differences were found in the display of social and conflict behaviors, nor in ERα expression patterns in the brain. We conclude that increases in serotonin levels during early development do not have long-term consequences for female sexual behavior in adulthood.

## 1. Introduction

Serotonin (5-HT) plays an important role in many social behaviors, including sexual behavior. While 5-HT is mainly a modulatory neurotransmitter during adulthood, 5-HT acts as a neurotrophic factor during early brain development, regulating cell division, differentiation, migration, and synaptogenesis (Azmitia, 2001; Gaspar et al., 2003). Therefore, it is assumed that changes in 5-HT levels during *in utero* neurodevelopment have the potential to affect these processes as well as subsequent serotonergic function (Lesch and Mossner, 1998). The occurrence of serotonin release during the different developmental stages, perinatally or in adulthood, could thus have different effects on behavioral outcomes.

Female sexual behavior is highly dependent on the estrous cycle, which is under regulation of the hypothalamic-pituitary-gonadal (HPG) axis. The principle of this axis is that the hypothalamus releases gonadotropin-releasing hormone (GnRH), which induces the pituitary to release follicle stimulating hormone (FSH) and luteal hormone (LH). The release of FSH and LH, in turn, stimulates the ovaries to release steroid hormones, like estrogen and progesterone (reviewed in (Snoeren, 2019)). Puberty arises when estrogens are secreted by the ovaries. During prepubertal development, neurotransmitter systems regulate the onset of puberty in the female rat through the modulation of GnRH and gonadotrophins (Moran et al., 2013; Rondina et al., 2003). There is close interaction between serotonin and gonadal hormones. The serotonergic system is regulated by sex steroids (James et al., 1989; Osterburg et al., 1987), and conversely, serotonin acts on the secretion of ovarian hormones (Jacobsen et al., 2015; Koppan et al., 2004; Tanaka et al., 1993; Terranova et al., 1990). Moreover, 5-HT has been detected in the ovaries, oviducts and uterus (Amenta et al., 1992). This suggests that the serotonin system is involved in the development of the HPG axis (Deneris and Gaspar, 2018; Millard et al., 2017). Since the activation and coordination of the HPG-axis is required for puberty establishment (Moran et al., 2013), it is then not surprising that repeated exposure to FLX *in utero* and during lactation was found to delay puberty onset in female Wistar rat offspring (Dos Santos et al., 2016) and increase the reproductive cycle length (Moore et al., 2015). Altogether, these findings indicate that 5-HT exposure during early development alters the estrous cycle in young animals, which could then change female sexual functioning in adulthood as well.

Serotonin can play a dual role in female sexual behavior via diverse receptor expression patterns in different brain regions (reviewed in (Snoeren et al., 2014)). Interestingly, 5-HT_1A_ receptor agonists strongly inhibit paracopulatory behavior (Snoeren et al., 2011a; Snoeren et al., 2011b) and lordosis responses in different strains of rats (Ahlenius et al., 1989; Kishitake and Yamanouchi, 2003; Mendelson and Gorzalka, 1986), whereas the effects of drugs that increase serotonin availability in the synapse, such as SSRIs, have shown conflicting results in female rats. For example, chronic fluoxetine treatment administered in adulthood inhibits female rat sexual activity in different strains (Adams et al., 2012; Matuszczyk et al., 1998; Uphouse et al., 2006), while paroxetine did not show any effects on lordosis or paracopulatory behaviors in Wistar female rats (Kaspersen and Ågmo, 2012; Snoeren et al., 2011b), possibly explained by desensitization of 5-HT_1A_ receptors (Snoeren et al., 2011b). Furthermore, lifelong increased 5-HT levels such as in serotonin transporter (SERT) knockout Wistar rats (Homberg et al., 2007), neither affects female sexual functioning (Snoeren et al., 2010). This illustrates how both the altered 5-HT signaling and the specific time window within life of such interventions contribute to effects on sexual functioning.

Although many studies have been performed on the role of serotonin on female rat sexual behavior in adulthood, not much is known about the role of serotonin release during early development on sexual functioning in adulthood. The few studies that investigated the effects of *perinatal* SSRI exposure on female sexual behavior are rather contradictive. Some studies showed a slight decrease in Wistar rat sexual behavior upon perinatal SSRI exposure (Houwing et al., 2019), while others suggested that postnatal fluoxetine exposure plays a stimulatory role on paracopulatory and receptive behaviors in Sprague-Dawley rats (Rayen et al., 2014). In addition, another study did not find any effect of early-life fluoxetine exposure on female Wistar rat sexual behavior (Dos Santos et al., 2016).

A limitation of the previous studies was the use of traditional test set-ups in which rats have a limited amount of both space and time to interact with each other. In addition, the use of only pairs of rats limits the opportunity to explore social interaction and does not model the natural situation in which rats live in groups (Calhoun, 1963; Robitaille and Bovet, 1976; Schweinfurth, 2020). In nature, rats copulate in groups consisting of one or several females and males (Calhoun, 1963; Robitaille and Bovet, 1976), and the mating patterns in groups of rats are quite different from the mating patterns observed in the traditional laboratory mating tests with pairs of rats or mice (Chu and Agmo, 2014, 2015b; Garey et al., 2002; McClintock, 1984; McClintock and Adler, 1978; McClintock and Anisko, 1982; McClintock et al., 1982; Snoeren et al., 2015).

The aim of this study was to investigate the effect of increased 5-HT levels during early development on female sexual behavior in adulthood. Mothers were treated with the SSRI fluoxetine during pregnancy and the lactation period. Since antidepressants can cross the placenta and are present in breast milk (Kristensen et al., 1999; Rampono et al., 2004), this procedure exposed the offspring to fluoxetine and increased the levels of serotonin during early development. The use of the seminatural environment does not only allow us to evaluate the effects of perinatal SSRI exposure on natural rat mating behavior, but also gives us the opportunity to study sexual functioning throughout the full behavioral estrus instead of during a short 30-minute copulation test (Chu and Agmo, 2014). As 5-HT is thought to affect the reproductive cycle, we wanted to investigate whether perinatal fluoxetine exposure disturbs female sexual behavior in adulthood via the estrogen receptor α (ERα). It was shown that postnatal clomipramine, a tricyclic antidepressant, increases the number of ERα-IR cells in the preoptic area, basolateral amygdala, medial amygdala, and septum (Limon-Morales et al., 2019; Molina-Jimenez et al., 2019), but reduces the number of ERα-positive cells in the hippocampal CA1/CA2 regions and raphe nucleus in male rats (Limon-Morales et al., 2019). The medial preoptic nucleus (MPOM), ventromedial nucleus of the hypothalamus (VMN), medial amygdala (MeA), bed nucleus of stria terminalis (BNST), and the dorsal raphé nucleus (DRN) are brain regions known for its role in the regulation in female rat sexual behavior (Arendash and Gorski, 1983; Hoshina et al., 1994; Kato and Sakuma, 2000; Kondo and Sakuma, 2005; Mathews and Edwards, 1977; Pfaff and Sakuma, 1979), and contain high levels of ERα (Laflamme et al., 1998; Yamada et al., 2009).We therefore also investigated the effects of perinatal fluoxetine exposure on ERα expression levels in these brain regions as potential working mechanism for a disturbed sexual behavior.

## 2. Material and Methods

The data was collected from video recordings obtained in a previously performed experiment. The materials and methods are therefore similar to those described previously (Heinla et al., 2020b; Houwing et al., 2019).

### 2.1 Animals and dam housing conditions

Male and female Wistar rats (n=10, weighing 200-250 g at the time of arrival, and obtained from Charles River (Sulzfeld, Germany)) were used for breeding. Before and after breeding, all animals were housed in same sex pairs in Makrolon® IV cages in a room with controlled temperature (21 ± 1 °C) and humidity (55 ± 10 %) on a 12:12 h light/dark cycle (lights on 11:00 AM). Commercial rat pellets (Standard chow from SDS, Special Diet Services) and tap water were provided ad libitum, and nesting material was provided.

All experimentation was carried out in agreement with the European Union council directive 2010/63/EU. The protocol was approved by the National Animal Research Authority in Norway.

### 2.2 Breeding and antidepressant treatment

To screen for their estrous cycle, dams were placed together with a male rat daily for a maximum of 5 minutes. When a lordosis response was observed during this period, they were considered in proestrus and thus ready for breeding. The breeding procedure was started by housing the receptive dam together with one male for approximately 24 hours (gestational day 0) in a Makrolon® IV cage. After the breeding, both male and female returned to their original homecage with a same-sex partner. The dams were pair-housed during pregnancy until gestational day 14, when the females were housed singly in Makrolon® IV cages with access to nesting material.

The antidepressant treatment was started on gestational day 1 (G1) and lasted until postnatal day 21 (PND21). The dams were administered daily with either 10 mg/kg fluoxetine (Apoteksproduksjon, Oslo, Norway) or a vehicle (Methylcellulose 1%, (Sigma, St. Louis, MO, USA)) using oral gavage with a stainless steel feeding needle (total of 6 weeks). The fluoxetine treatment was prepared with tablets (for human usage) that were pulverized and dissolved in sterile water (2mg/mL) and injected at a volume of 5mL/kg. Methylcellulose powder, the non-active filling of a fluoxetine tablet, was used as control condition. The powder was dissolved in sterile water to create a 1% solution and administered at a volume of 5mL/kg as well. The weight of each female was measured every third day and the amount of vehicle/fluoxetine treatment adjusted accordingly. The dose of fluoxetine was based on previous literature comparing fluoxetine blood levels of humans and animal (Lundmark et al., 2001; Olivier et al., 2011). The same dosage was used in previous experiments (Heinla et al., 2020a; Houwing et al., 2020a; Houwing et al., 2019; Houwing et al., 2020b). Near the end of pregnancy, dams were checked twice per day (9:00 h and 15:00 h) for pup delivery. The offspring were thus exposed to perinatal fluoxetine via the treatment of the dams (in uterus and via breast feeding).

### 2.3 Offspring housing conditions before the seminatural environment

After birth, litters were not culled. On postnatal day 21 (PND21), offspring pups were weaned, ears were punched for individual recognition, and they were housed in groups of two or three same sex littermates in Makrolon IV cages. The offspring were left undisturbed and were only handled during weekly cage cleaning, until introduction into the seminatural environment (at 13-18 weeks of age. (For more details on which rats were group-housed and belonged to each cohort of the seminatural environment, see supplemental materials of (Houwing et al., 2019))

### 2.4 Ovariectomy surgery and hormone treatment

Two weeks before the start of the experiment, female offspring were ovariectomized under isoflurane anesthesia, and pre-surgery analgesia of buprenorphine (.05 mg/kg, subcutaneously) and Carprofen (5mg/kg, subcutaneously). The females were placed on their ventral surface, and a 1-2 cm longitudinal midline dorsal skin incision was made at the lower back of the animal. Via bilateral incisions in the muscles, the ovaries were located in the peritoneal cavity and extirpated. The muscle incisions were sutured and a wound clip was placed for skin closure. Post-surgery analgesia consisted of Carprofen (5mg/kg subcutaneously) 24 and 48 hours after surgery. Female offspring were single-housed for 3 days during recovery before returning to their home cage.

In order to induce receptivity in all females on the same day of observation (day 7) in the seminatural environment, the females received 18 µg/kg estradiol benzoate on day 5, and 1 mg of progesterone on day 7 at 10:00 am. Estradiol benzoate and progesterone (Sigma, St Louis, MO, USA) were dissolved in peanut oil (Apoteksproduksjon, Oslo, Norway) and injected in a volume of 1 ml/kg. This hormonal treatment is known to induce similar hormonal patterns as those in intact females (Fadem et al., 1979)

### 2.5 Seminatural environment

The seminatural environment (2.4 x 2.1 x 0.75 meters) consisted of a burrow system and an open field area, which were connected by four 8 x 8 cm openings (a figure of the environment can be found in (Chu and Agmo, 2014; Houwing et al., 2019; Snoeren et al., 2015). The burrow system contained several tunnels (7.6 cm wide and 8 cm high) and four nest boxes (20 x 20 x 20 cm), and was covered with Plexiglas. The open area, on the other hand, was an open space with two partitions (40 x 75 cm) to create obstacles simulating nature. With walls of 75 cm, the open area was not covered with Plexiglas. A curtain between the open area and the burrow system allowed the light intensity for both arenas to be controlled separately: total darkness for the complete day in the burrow area, and a simulated day-night cycle in the open area. The day-night cycle was obtained with the use of a lamp 2.5 m above the center of the open area provided light (180 lux) from 10:45 pm to 10:30 am (simulating day light). From 10:30 am to 11:00 am the light intensity gradually decreased to approximately 1 lux, the equivalent of full moonlight. Similarly, the light gradually increased again from 1 to 180 lux from 10:15 pm to 10:45 pm.

Rats could freely walk from and to the open area and the burrow system. A 2 cm layer of aspen wood chip bedding (Tapvei, Harjumaa, Estonia) was covering the floor of both the open area and the burrow system. In addition, as environmental objects, the nest boxes contained 6 squares of nesting material each (nonwoven hemp fibers, 5 x 5 cm, 0.5 cm thick, Datesend, Manchester, UK), and 3 red polycarbonate shelters (15 x 16.5 x 8.5 cm, Datesend, Manchester, UK) and 12 aspen wooden sticks (2 x 2 x 10 cm, Tapvei, Harjumaa, Estonia) were randomly distributed within the open area. Food was provided in one big pile of approximately 2 kg, in front of the open area wall opposite of the openings. Water was available *ad libitum* in four water bottles located in the lower right corner of the open field.

Video cameras were mounted on the ceiling 2 m above the seminatural environment: one above the open field (Basler) and an infrared video camera above the burrow system (Basler). Videos were recorded using Media Recorder 2.5. Cameras were connected to a computer and data was (immediately) stored on an external hard drive. Every 24 h, the recording was manually stopped and restarted to create recordings with a length of 24h. This was done to make sure that if a recording error should occur during the 8 day period, only one recording day would be lost.

### 2.6 Procedure

The day before introduction to the seminatural environment, offspring were shaved and marked under isoflurane anesthesia for individual recognition on video (the wound clips of the females were also removed at the same time). For more details, see (Houwing et al., 2019).

At 10 am on the first day of the experiment (day 0), a cohort of 4 male and 4 female offspring (from different litters) was placed in the seminatural environment for 8 days. Each cohort contained 2 CTR-females, 2 CTR-males, 2 FLX-females, and 2 FLX-males. In total, 5 cohorts of rats entered the seminatural environment, resulting in an experimental number of 10 rats for each treatment per sex. All rats were sexually naïve when entering the seminatural environment.

On day 5 and 7 at 10: am, the females were injected with estradiol benzoate and progesterone respectively. A trained observer screened the video recordings on day 7 after the progesterone treatment, and searched for the first displayed lordosis for each female. This was considered the onset of the *behavioral estrous period*. After this point, all social and sexual behaviors mentioned in table 1 were scored until the last lordosis was observed (the end of the behavioral estrous period). In order to only analyze the periods in which the females are sexually active, we divided the behavioral estrous period into *copulation bouts*. One copulation bout started with the first lordosis of the bout, and ended with the last lordosis after which the female does not show another lordosis for at least one hour. After this hour, a new lordosis is the start of the next copulatory bout (see Figure 1). The copulation bout that contained most lordosis responses was considered the *most active bout*.

**Table 1.** Ethogram of observed female behaviors in the seminatural environment

| **A) Behavior** | **Description** |
| --- | --- |
| **Lordosis** | Receptive behavior with a hollow back and deflect of tail to one side |
| **Paracopulatory behavior** | Female approaching a male followed by runaway, often associated with darts, hops, ear wiggling |
| **Mount (received)** | Mounting on the rump of the female from behind with pelvic thrusting (received) |
| **Intromission (received)** | Mount including penile insertion (received) |
| **Ejaculation (received)** | Penile insertion that lasts longer than an intromission and is associated with rhythmic contractions (received) |
| **Sniffing others** | Sniffing any part of the conspecifics body, except for the anogenital region |
| **Sniffing anogenitally** | Sniffing the anogenital region of the conspecific |
| **Pursuing/Chasing** | Moving or running forward in the direction of a conspecific |
| **Boxing/Wrestling** | One or both animals are pushing, pawing and grabbing at each other using their forepaws |
| **Nose-off** | Facing another rat, usually in a tunnel, resulting in one rat moving forwards and the other backing up |
| **Fighting** | Forming a tight ball with another rat, rolling around while biting |
| **Rejection** | Rejecting another rat by a leg kick or box |
| **Any other behavior** | All the behaviors that do not fit any of the above mentioned categories |
| **B) Behavioral clusters of observed behaviors in the seminatural environment** |  |
| **Total copulations** | Total number of received mounts, intromissions and ejaculations |
| **Social behavior** | Combines sniffing others, sniffing anogenitally, and pursuing/chasing (previously called *active social behavior* (Houwing et al., 2019)) |
| **Conflict behavior** | Combines boxing/wrestling, nose-off, fighting, and rejection |

**Figure 1:**
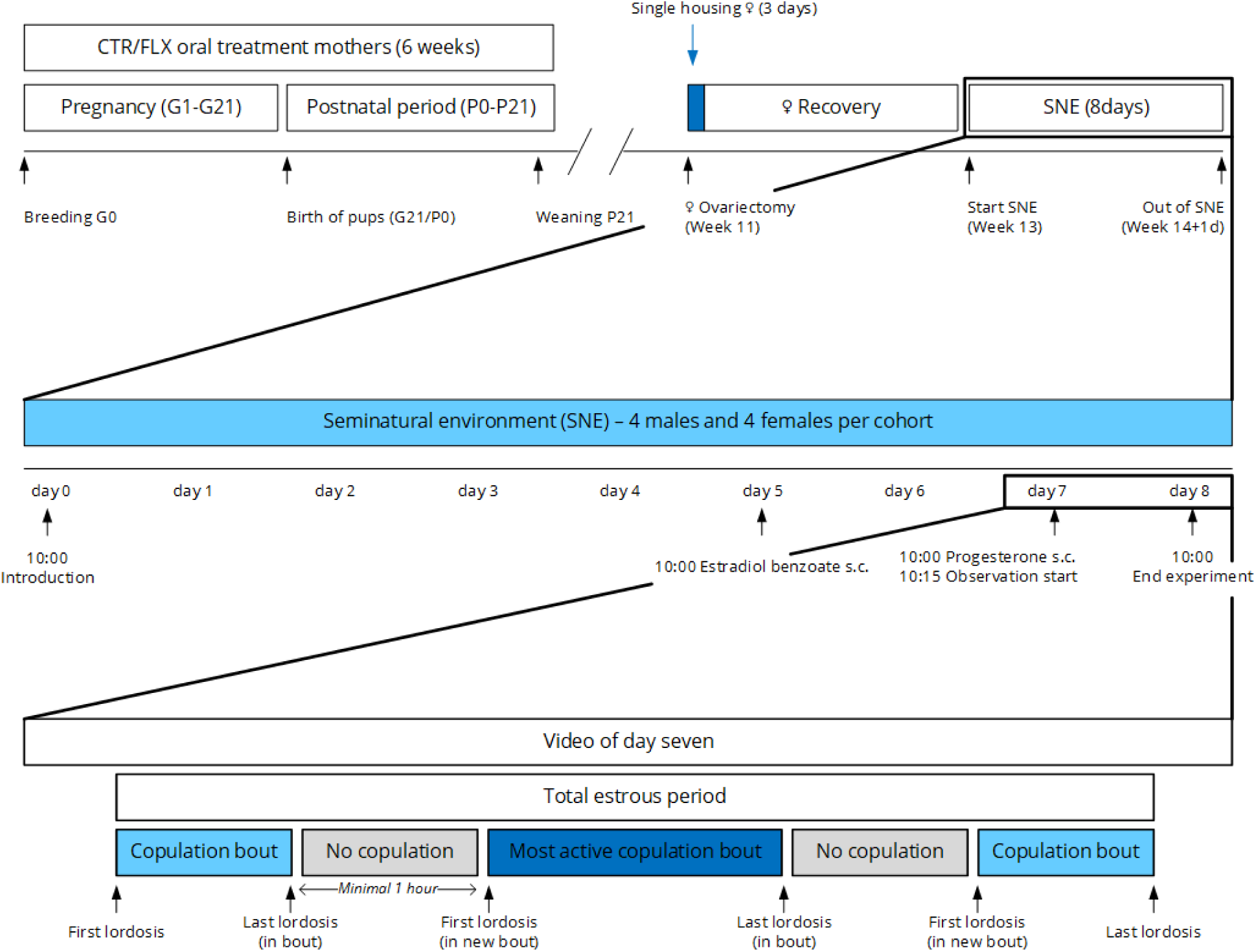
Schematic overview of all experimental procedures. CTR = control, FLX = fluoxetine, G = gestational day, PND =postnatal day, SNE = seminatural environment.

The frequencies and durations of a wide variety of behaviors during all copulation bouts within the behavioral estrous period were scored by an observer blind for the treatment of the animals (Table 1). In addition, the location of the animal was scored: in the open field or in the burrow. During interactions with other animals, the interacting partner was also noted. All behavioral scoring was done using the Observer XT, version 12 (Noldus, Wageningen, the Netherlands).

### 2.7 ERα expression patterns with immunohistochemistry

To investigate the effects of perinatal fluoxetine exposure on the estrogen receptor alpha (ERα) expression in the brain, immediately after exiting the seminatural environment, the rats were euthanized with an overdose of pentobarbital. Subsequently, they were perfused transcardially with PBS followed by 4% formaldehyde. The brain was removed and postfixed overnight at 4°C in 4% formaldehyde. It was then rinsed and cryoprotected in (sequentially) 10%, 20%, and 30% sucrose in PBS. After cryoprotection, the brains were snap frozen in isopentane cooled on dry ice, and stored in glas vials at −80°C until further processing.

The brains were frozen-sectioned in 40µm with a sledge microtome. Sections containing the brain regions of interest were collected and processed in accordance with a conventional free-floating protocol. The brain regions of interest, of which the coordinates were determined using the rat brain atlas Paxinos&Watson (6th edition), were the medial preoptic nucleus (MPOM, anteroposterior (AP) between −0.12 and −0.96), the ventrolateral part of the ventromedial nucleus of the hypothalamus (VMHvl, AP between −2.2 and −3.1), the posterodorsal part of the medial amygdala (MePD AP between −2.9 and −3.4), the bed nucleus of the stria terminalis (BNST, AP between −0.4 and −0.96) and the dorsal (DRD) and ventral (DRV) part of the dorsal raphe nucleus (DRN, AP between −7.2 and −7.9).

For immunohistochemistry, sections were washed in 0.1M Tris-buffered-saline (TBS), blocked for 30 min in 0.5% BSA, and incubated on an orbital shaker for 24h at room temperature + 72h at 4 °C in primary antibody (polyclonal rabbit anti-ERα, Merck-Millipore, cat. 06-935, dilutions: MPOM/BNST 1:10 000, VMN/MePD 1:5 000, and DRD/DRV 1:3 000) solution containing 0.1% Triton-X and 0.1% BSA in TBS. Sections were then incubated in secondary antibody (biotinylated goat anti-rabbit, Abcam, cat. ab670, dilution: 1:400) solution containing 0.1% BSA in TBS for 30 min., avidin-biotin-peroxidase complex (VECTASTAIN ABC-HRP kit, Vector laboratories, cat. PK-6100, dilution: 1 drop A + 1 drop B in 10 mL TBS) solution for 30 min., and 3,3’-diaminobenzidine solution (DAB substrate kit (HRP), Vector laboratories, cat. SK-4100, dilution: 1 drop R1 + 2 drops R2 + 1 drop R3 in 5 mL water) for 5 min., with TBS washes in between all steps. This resulted in brown staining of ERα-containing cells. After coverslipping, the slides were loaded into an Olympus VS120 virtual slide microscope system. Image scans were obtained for each section using a 20x objective (NA 0.75), automatic focus settings in single plane, and fixed exposure of 1.13 ms (MPOM/BNST) and 1.11 ms (VMN/MePD and DRD/DRV). Using Olyvia online database software, high resolution cropped images (5x digital zoom) of the brain regions of interest were downloaded and opened in Corel PHOTO-PAINT 2017, and a zone of always the same size and location was drawn to fit into the regions of interest. The following sizes of the zones were used: MPOM 0.333×0.558 mm, VMNvl 0.222×0.222 mm, MePD 0.278×0.5 mm, BNST 0.5×0.278 mm, DRD and DRV 0.222×0.278 mm.

On these photomicrographs, all stained cells inside the zones were manually counted using Cell Counter plugin in ImageJ. This was performed for three sections per brain area per rat (except for the BNST, where 6 rats only resulted in 2 sections per rat). Each section was counted twice when less than 5% difference in cell count was obtained, and 3 times when more than 5% difference was observed. Then the average of the two counts with less than 5% count difference was taken as data point. Per rat and brain region, the average of the 3 sections was used for further analysis. The number of counted cells was divided by the surface area of the zone to obtain the outcome measure of number of ERα+ cells per mm^2^.

### 2.8 Data preparation and statistical analysis

As indicated in table 1, behavioral clusters were created beforehand by grouping relevant behaviors. In addition, the lordosis quotient was calculated for each female by dividing the total number of lordosis responses by the total number of received copulations x 100. This was done separately for each copulatory bout, in addition to the total behavioral estrous period (combining all copulatory bouts excluding the behaviors during the intermitting periods of sexual inactivity).

In addition, we controlled for the potential effects caused by the differences in lengths of the bouts and behavioral estrus. Therefore, all the behavioral parameters were divided by the duration of the bout or behavioral estrus in minutes.

In order to analyze the patterns of sociosexual behavior over the full course of the copulatory bouts, we divided the total duration of each bout into 5% timebins. Then we analyzed the behaviors performed in each 5% timebin and created cumulative data points over the full course of the bout. A similar analysis was performed on the behavioral estrous period. This was done on both the original parameters and the parameters controlled for the duration of the bouts.

At last, we analyzed all the above for the behaviors and behavioral clusters that were performed in the total environment, but also separate for the open field and the burrow. In addition, we determined each female’s preferred male (towards who they showed most lordosis responses), and compared the sociosexual behavior towards the preferred male and the non-preferred males.

A Shapiro–Wilk test showed homogeneity of variance in all behaviors, except for the data controlled for the length of the bouts. The behavioral data that was normally distributed, was therefore analyzed using the independent *t*-test to compare FLX-females with CTR-females, while the data per minute was analyzed with the nonparametric Mann–Whitney U test. A repeated measures ANOVA was used for determining whether there were significant changes in a variable over the 20 5% intervals of the copulatory bouts of the behavioral estrous period. In case of significance, a bonferroni post-hoc analysis was made for different intervals in order to detect at which point during estrus the differences between CTR- and FLX-females occurred. No effects of litter were found.

## 3. Results

As mentioned in the methods, the behavioral estruses of the female rats were divided in multiple copulation bouts. Figure 2A shows a representation of these copulation bouts within the behavioral estrus, starting after the injection of progesterone. The copulation bout in which the female performed most lordoses was considered the *most active bout*. No differences were found between the duration of the behavioral estrus (*t*_(18)_=0.077, NS, Figure 2B), the onset of the *most active bout* after progesterone injection (*t*_(18)_=-1.079, NS, Figure 2C), or the duration of the *most active bout* (*t*_(12.458)_=0.353, NS, Figure 2D) between CTR- and FLX-females.

**Figure 2:**
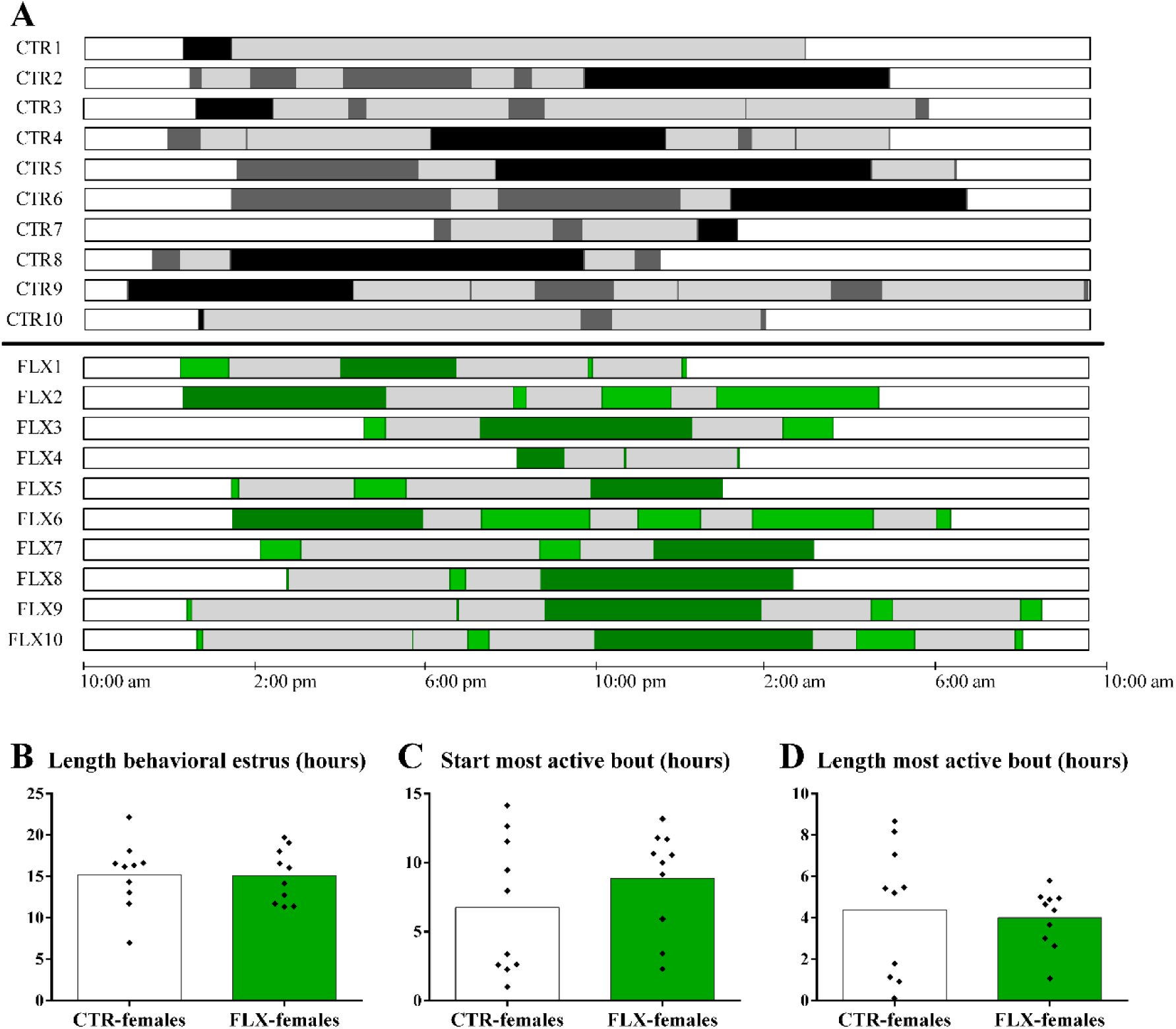
Overview of behavioral estrus divided in copulation bouts. A) Schematic overview of the timing, length and number of different copulation bouts. The intermitting periods of sexual inactivity are shown in light grey and the copulation bouts in dark grey (CTR-females, upper part) or light green (FLX-females, lower part). The most active bouts are shown in black (CTR-females) and dark green (FLX-females). The behavioral estrus starts with the first lordosis and thus with the onset of the first copulation bout, and ends with the last lordosis and thus at the end of the last copulation bout, B) the length of the total duration of the behavioral estrus in hours, C) the onset of the most active bout in hours after progesterone injection (at 10:00 am), D) the length of the total duration of the “most active bout” in hours. Data are shown with individual data points, with the bars representing the mean.

### 3.1 Sexual behavior

When the data of the full behavioral estrus was analyzed, no differences were found in the time spent on paracopulatory behavior (*t*_(18)_=-0.558, NS, Figure 3A) or the number of paracopulatory behavior episodes (*t*_(18)_=-0.821, NS, Table S1) between CTR- and FLX-females. When the pattern of darts and hops was analyzed over the time course of the behavioral estrus (in 5% time bins), FLX-females showed no significant difference in pattern of time spent on (timebin x treatment: F_(19,342)_=0.899, NS, Figure 3B) and number of paracopulatory behaviors (timebin x treatment: F_(19,342)_=1.055, NS, data not shown) as CTR-females. In addition, FLX-females showed no differences in lordosis quotient as CTR-females (*t*_(18)_=0.893, NS, Figure 3C), just as the number of lordoses response over the time course of the behavioral estrus were not significantly different between CTR- and FLX-females (timebin x treatment: F_(19,342)_=0.453, NS, Figure 3D). It is therefore not surprising that there was no significant difference between CTR- and FLX-females in number of received copulations (*t*_(18)_=0.598, NS, Figure 3E, mounts, intromissions and ejaculations are shown separately in Table S1). Again, when the pattern of received copulations was analyzed over the time course of the behavioral estrus, no differences in the received copulatory behavior pattern was found for FLX-females compared to CTR-females (timebin x treatment: F_(19,342)_=0.530, NS, Figure 3F). Only when the number of mounts and intromissions were analyzed separately, it was found that the FLX-females received less mounts (timebin x treatment: F_(19,342)_=2.356, p=0.001) and intromissions (timebin x treatment: F_(19,342)_=2.070, p=0.006) in the burrow area than CTR-females in the 30% to 70% timebins, but these mounts and intromissions were then partly received in the open area of the seminatural environment instead of in the burrow (Figure S1).

**Figure 3:**
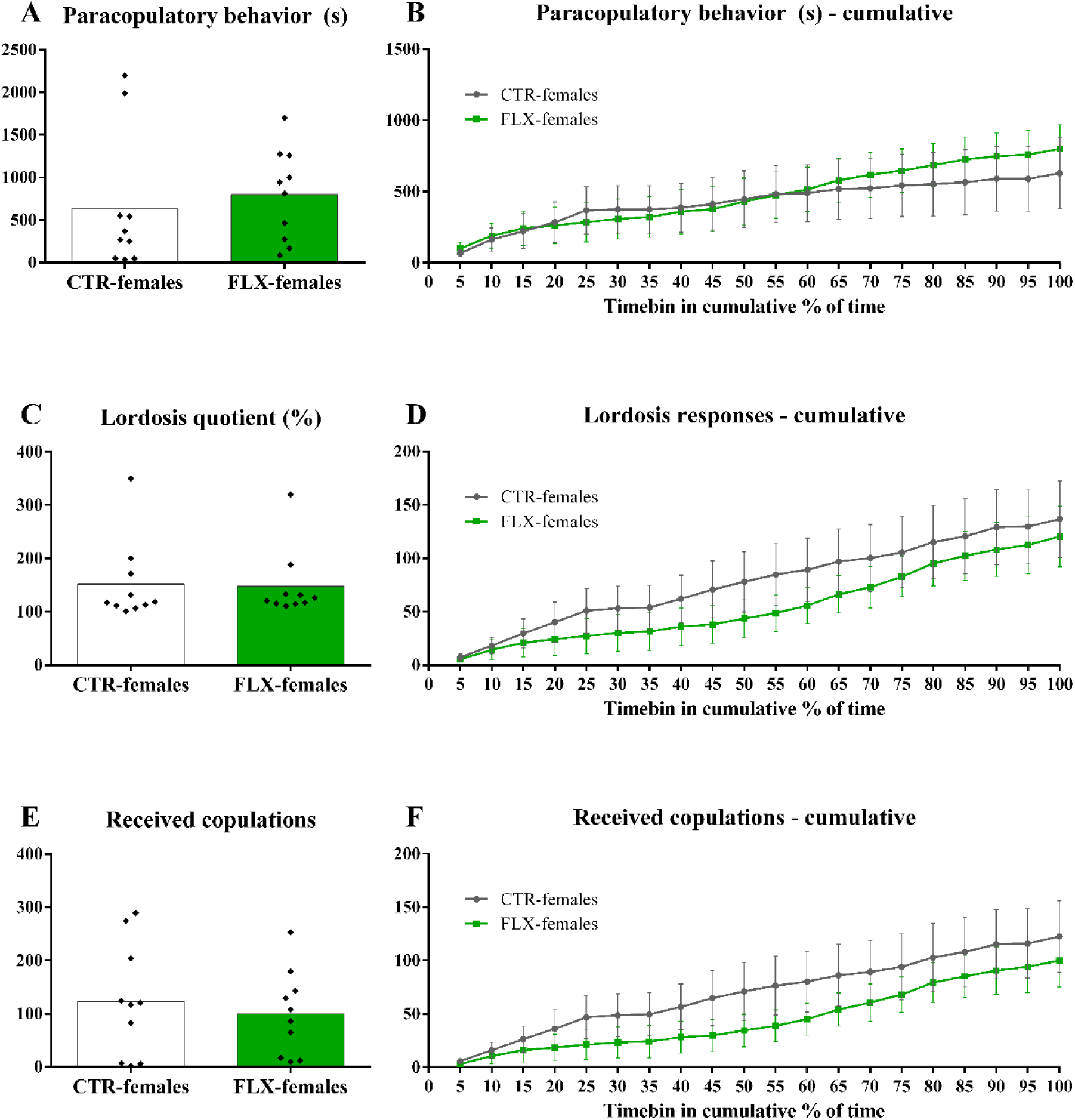
Sexual behaviors during the full behavioral estrus. The data represents A) the time spent on paracopulatory behaviors in seconds, B) the time spent on paracopulatory behaviors in seconds per cumulative 5% timebin of the behavioral estrus, C) the lordosis quotient (lordosis responses/received copulations * 100%), D) the number of lordoses responses per cumulative 5% timebin of the behavioral estrus, E) the number of received copulations (mounts+intromissions+ejaculations), and F) the number of received copulations per cumulative 5% timebin of the behavioral estrus. On the left, the data are shown with individual data points, with the bars representing the mean. On the right, the data are shown as mean±sem for CTR- and FLX-females.

Since female rats perform sexual behavior in multiple copulation bouts, we also analyzed the behaviors performed in the bout during which the female showed most lordosis responses: the *most active bout*. However, no differences in sexual behavior were found between CTR- and FLX-females in the *most active bout* compared to the full behavioral estrus. As shown in Figure 4, no differences were found between CTR- and FLX-females in the amount of time spent on paracopulatory behavior in the *most active bout* (*t*_(18)_=0.766, NS, Figure 4A), numbers of paracopulatory behavior episodes (*t*_(18)_=0.539, NS, Table S1), lordosis quotients (*t*_(18)_=0.863, NS, Figure 4C), and number of received copulatory behaviors (*t*_(18)_=0.626, NS, Figure 4E). Also when the behavioral patterns over the time course of the *most active bout* were analyzed, no differences in the time spent on paracopulatory behavior (timebin x treatment: F_(19,342)_=0.242, NS, Figure 4B), the number of lordosis responses (timebin x treatment: F_(19,342)_=0.151, NS, Figure 4D), or the received copulations (timebin x treatment: F_(19,342)_=0.172, NS, Figure 4F) were found between CTR- and FLX-females. The lower amounts of received mounts and intromissions in the burrow area (instead of open area) was no longer significantly different between CTR- and FLX-females during the *most active bout*.

**Figure 4:**
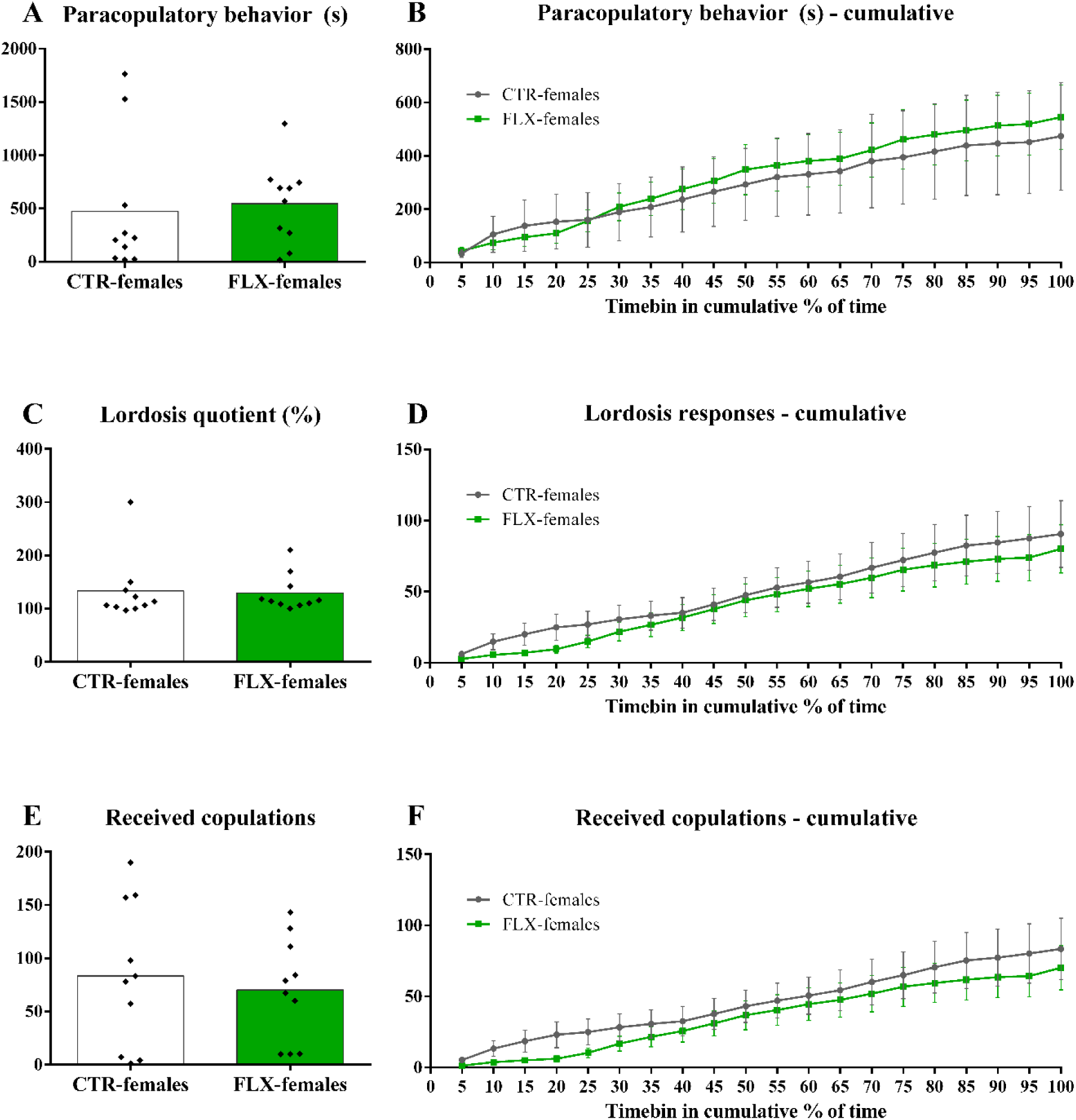
Sexual behaviors during the most active bout. The data represents A) the time spent on paracopulatory behaviors in seconds, B) the time spent on paracopulatory behaviors in seconds per cumulative 5% timebin of the “most active bout”, C) the lordosis quotient (lordosis responses/received copulations * 100%), D) the number of lordoses responses per cumulative 5% timebin of the most active bout, E) the number of received copulations (mounts+intromissions+ejaculations), and F) the number of received copulations per cumulative 5% timebin of the most active bout. On the left, the data are shown with individual data points, with the bars representing the mean. On the right, the data are shown as mean±sem for CTR- and FLX-females.

We also determined the preferred male per female, which was the male who elicited the most lordoses responses. Then we looked at whether CTR- and FLX-females showed any differences in sexual behavior towards the preferred or non-preferred males. The data revealed that there were also no significant differences in sexual behaviors between CTR- and FLX-females towards the preferred or non-preferred males (data not shown).

Since the duration of both the behavioral estrus and the *most active bout* are different between all females, they could theoretically have had more or less time to perform the sexual behaviors. Although FLX-females had on average the same lengths of the behavioral estrus and *most active bout* as CTR-females, we still performed an analysis in which we controlled for these differences in bout durations by dividing the behaviors by the amount of minutes in the behavioral estrus or *most active bout*. The analysis of this data revealed no relevant effects, and therefore the results will not be described in detail. Only in the time spent on paracopulatory behavior (in seconds per minute) over the time course of the full behavioral estrus there was significant interaction effect of timebin and treatment between CTR-and FLX-females (timebin x treatment: F_(19,342)_=2.974, p<0.01, Fig S2), but this effect was not confirmed with posthoc analysis. In addition, the data revealed a significant different pattern (a decrease) of received copulations (per minute) of FLX-females in the burrow area during the full behavioral estrus (timebin x treatment: F_(19,342)_=2.582, p<0.01), which was again compensated in the open area as the number of received copulations (per minute) in the whole seminatural environment did not differ from CTR-females. During the course of the *most active bout*, a decrease in number of received mounts (per minute) was again found (timebin x treatment: F_(19,342)_=1.777, p=0.021) as mentioned before. During this bout, however, there was no effect on the total number of received copulations (Figure S3).

### 3.2 Social behaviors

Sexual behavior can be dependent on social and conflict behavior. The use of a seminatural environment allows for observations of these behaviors as well. We have therefore also scored the social behaviors consisting of sniffing others, anogenitally sniffing, allogrooming, and pursuing/chasing other rats, and conflict behaviors entailing fighting, boxing/wrestling, nose-off, and rejections. When these behaviors were analyzed over the full course of the full behavioral estrus, it was found that one CTR-female showed an extreme amount of sniffing behavior. Although the data-analysis with and without this female did not differ, the data presented below is analyzed without this outlier.

Data analysis revealed that FLX-females spent no significant difference in time on social behaviors as CTR-females (*t*_(17)_=0.416, NS, Figure 5A). In addition, they showed no difference in number of social behavior episodes (*t*_(17)_=0.633, NS, Table S1). When the social behavior was analyzed over the course of full behavioral estrus, it was found that FLX-females show a significantly different pattern in time spent on social behavior than CTR-females (timebin x treatment: F_(19, 323)_=2.304, p=0.02, Figure 5B), but this effect was not confirmed in the posthoc analysis of the different 5% timebins. It should be mentioned as well that this effect does not persist when the outlier CTR-female was included in the analysis. FLX-females also did not show a different pattern in the number of social behavior episodes over the time course of the behavioral estrus (frequency: timebin x treatment: F_(19,323)_=0.997, NS). In terms of conflict behavior, there were no significant differences found in how long and often FLX-females were involved in conflict situations compared to CTR-females (duration: *t*_(18)_=0.668, NS, Figure 5C; frequency: *t*_(18)_=0.670, NS, Table S1). When this behavior was analyzed over the full time course of the behavioral estrus, FLX-females seemed to spent slightly less time in conflict behavior, but this difference was not significant (duration: timebin x treatment: F_(19,342)_=0.507, NS, Figure 5D; frequency: timebin x treatment: F_(19,342)_=0.510, NS). A posthoc analysis only showed a significant difference in the 10% timebin between CTR- and FLX-females.

**Figure 5:**
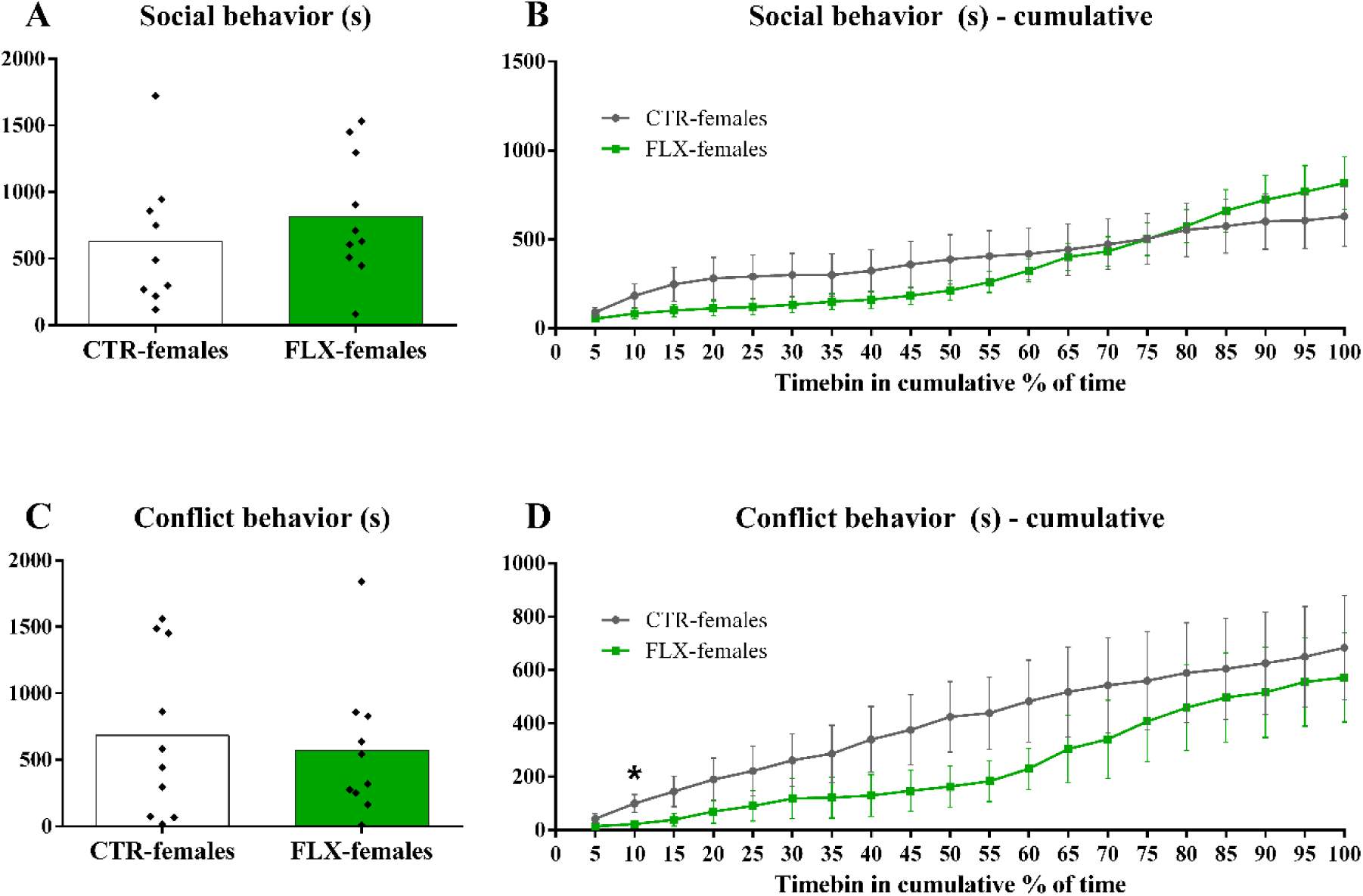
Social behaviors during the full behavioral estrus. The data represents A) the time spent on social behaviors in seconds, B) the time spent on social behaviors in seconds per cumulative 5% timebin of the behavioral estrus, C) the time spent on conflict behaviors in seconds, D) the time spent on conflict behaviors in seconds per cumulative 5% timebin of the behavioral estrus. On the left, the data are shown with individual data points, with the bars representing the mean. On the right, the data are shown as mean±sem for CTR- and FLX-females. * p<0.05 compared to CTR-females.

Also in terms of social or conflict behavior towards the preferred or non-preferred males, there were no differences between CTR- and FLX-females (data not shown).

When we analyzed the social behaviors during the *most active bout*, we did not find any relevant differences between CTR- and FLX-females. FLX-females spent the same amount of time on social behavior (*t*_(17)_=0.348, NS, Figure 6A) and conflict behavior (*t*_(18)_=0.969, NS, Figure 6C) as CTR-females, just as no differences were found in the number of social behavior (*t*_(17)_=0.667, NS, Table S1) and conflict episodes (*t*_(18)_=0.797, NS, Table S1). In addition, when the data was analyzed over the full time course of the *most active bout*, no differences in patterns in social behavior (duration: timebin x treatment: F_(19,323)_=0.924, NS, Figure 6C; frequency: timebin x treatment: F_(19,323)_=0.042, NS) and conflict behavior (duration: timebin x treatment: F_(19,342)_=0.168, NS, Figure 6D; frequency: timebin x treatment: F_(19,342)_=0.301, NS) were found between CTR- and FLX-females. A posthoc analysis of the social behavior did only find that FLX-females spent less time on social behavior during the 15% and 20% timebins.

**Figure 6:**
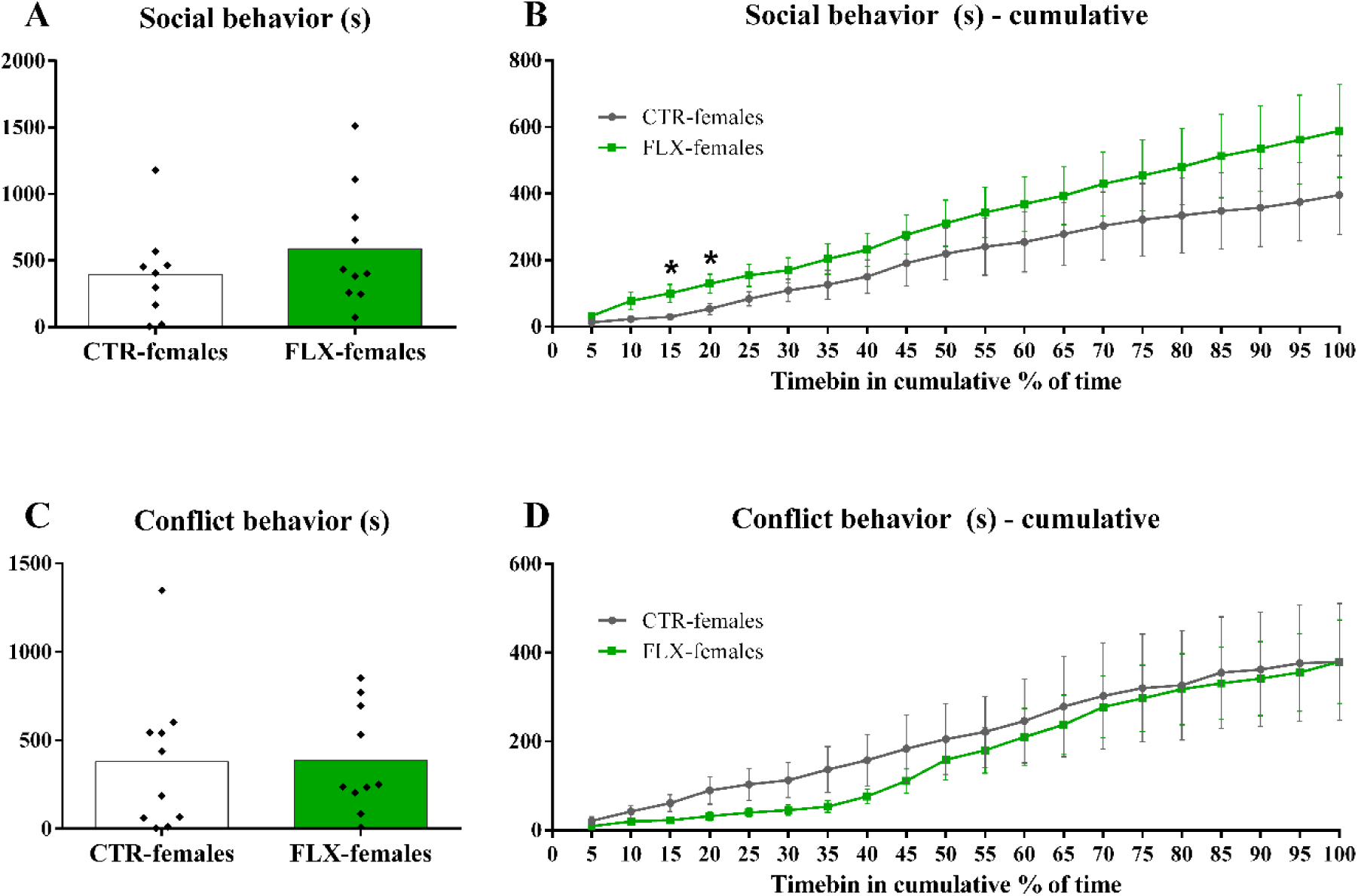
Social behaviors during the most active bout. The data represents A) the time spent on social behaviors in seconds, B) the time spent on social behaviors in seconds per cumulative 5% timebin of the most active bout, C) the time spent on conflict behaviors in seconds, D) the time spent on conflict behaviors in seconds per cumulative 5% timebin of the most active bout. On the left, the data are shown with individual data points, with the bars representing the mean. On the right, the data are shown as mean±sem for CTR- and FLX-females. * p<0.05 compared to CTR-females.

Again, an analysis that controlled for the duration of the behavioral estrus and *most active bout* was performed (Figure S4 and S5). While the small effects on active social and conflict behavior that were mentioned above disappeared, a significant change in pattern of social behavior over the time course of the full behavioral estrus was found between CTR- and FLX-females (timebin x treatment: F_(19,323)_=2.974, p<0.01). However, posthoc analysis did not confirm this finding.

### 3.3 Estrogen receptor α expression

After performing immunohistochemistry staining for the estrogen receptor α (ERα) on 19 of the 20 brains (due to practical issues we lost one brain), the number of ERα-containing cells were counted in the medial preoptic area (MPOM), the ventrolateral part of the ventromedial nucleus of the hypothalamus (VMNvl), the posterodorsal part of the medial amygdala (MePD), the bed nucleus of the stria terminalis (BNST) and the dorsal (DRD) and ventral (DRV) part of the dorsal raphe nucleus (DRN). As shown in Figure 7, no differences were found in the number of ERα-containing cells in the MPOM (*t*_(57)_=-1.168, NS), VMNvl (*t*_(55)_=0.565, NS), MePD (*t*_(55)_=1.001, NS), BNST (*t*_(48)_=0.884, NS), DRD (*t*_(52)_=0.875, NS), DRV (*t*_(53)_=-1.159, NS), and DRN (*t*_(53)_=-0.126, NS) between CTR- and FLX-females. A separate analysis of the more anterior and posterior parts of these brain regions resulted in similar ERα expression between FLX and CTR-females.

**Figure 7:**
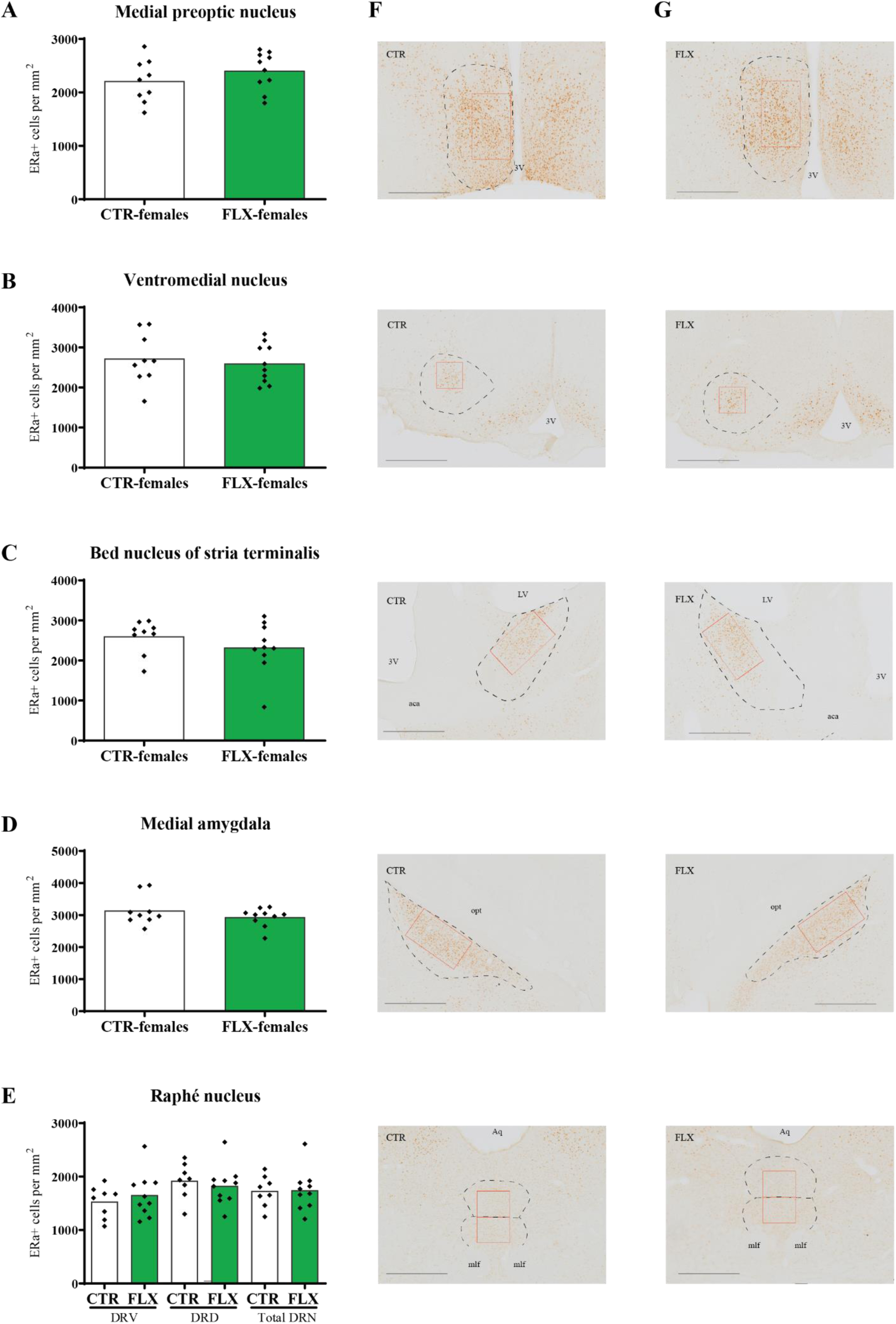
Estrogen receptor α containing cells. The data represents the number of ERα+ cells per mm^2^ in A) the medial preoptic nucleus, B) the ventromedial nucleus of the hypothalamus, C) the bed nucleus of stria terminalis, D) medial amygdala, and E) the dorsal raphé nucleus. The data are shown with individual data points, with the bars representing the mean for CTR- and FLX-females. On the right, an example of a microphotographs with the counted ERα+ cells of a F) CTR-female and G) FLX-female were shown per brain region. The red outlines represent the zone that was used for counting.3V = 3^rd^ ventricle, LV = lateral ventricle, ac = anterior commissure, opt =optic tract, Aq =Aqueduct, mlf = medial longitudinal fasciculus. The scale-line represents 0.5 mm.

## 4. Discussion

The aim of this study was to investigate the effect of perinatal SSRI exposure on female rat sexual behavior in adulthood. We evaluated the female sexual patterns during the full behavioral estrus from the first lordosis to the last, and found that female rats that were perinatally exposed to fluoxetine showed similar levels of sexual activity as control females. As summarized in Table 2, no differences were found in the time spent on and the number of paracopulatory and receptive behaviors between CTR- and FLX-females, nor did the patterns of these behaviors differ over the time course of the behavioral estrus. When we studied only the *most active bout*, the copulation bout during which the females performed most lordoses, again no differences in female sexual behavior were observed. Therefore, we conclude that the exposure to elevated serotonin levels during early development does not affect female sexual behavior in adulthood.

**Table 2.**
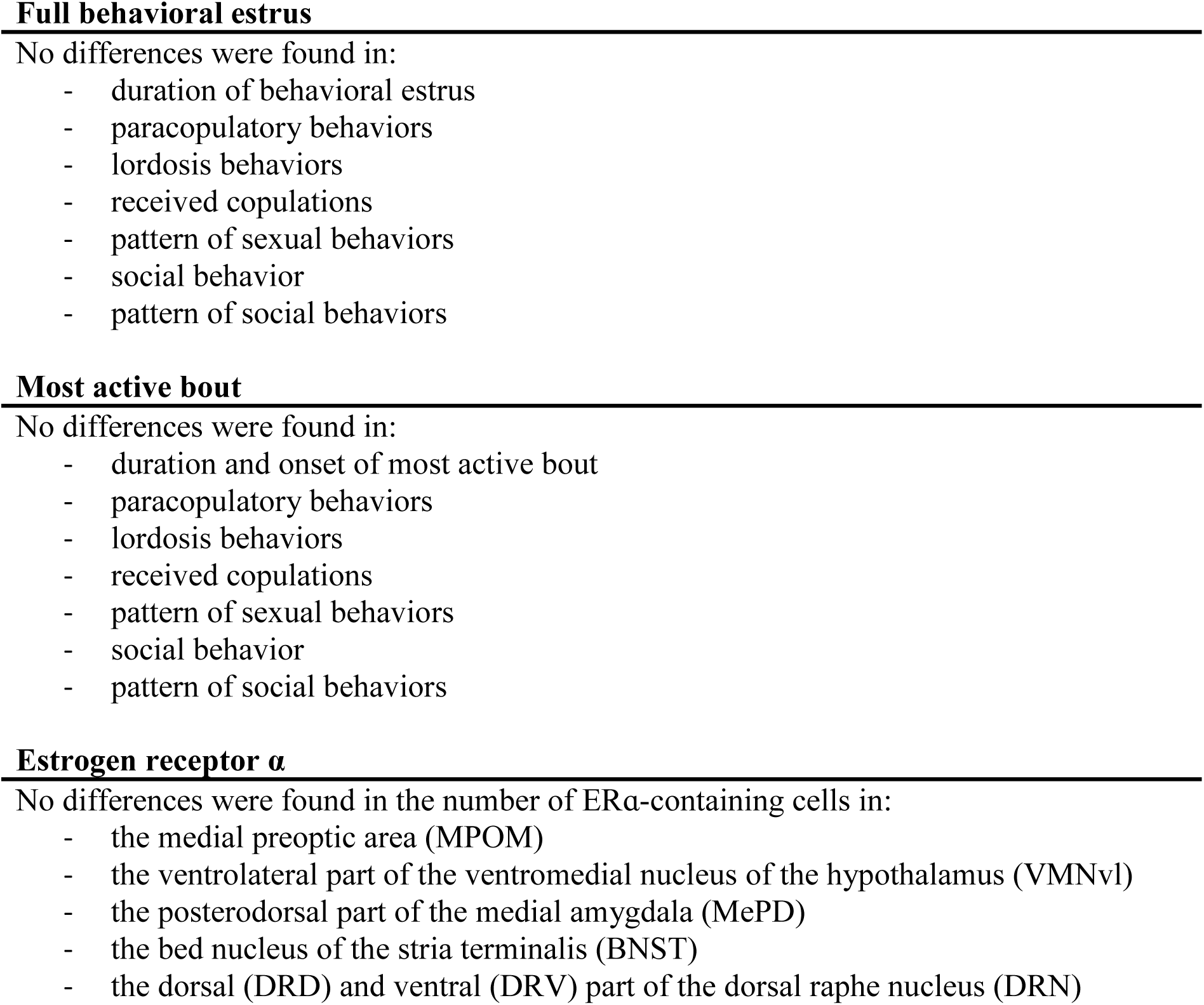
Summary of the findings

These findings were in line with another study in which perinatal FLX exposure did not affect lordosis behavior in naturally cycling adult female Wistar rats (Dos Santos et al., 2016). Another study, on the other hand, found that postnatal exposure to FLX facilitated paracopulatory and receptive behaviors in intact female Sprague-Dawley rats (Rayen et al., 2014). There are several dissimilarities in experimental design that could explain the different findings. First, (Rayen et al., 2014) primed the females with a high dose of estradiol benzoate (EB) 36 hours before behavioral testing. Although this method has been used previously to cause females to enter behavioral estrus (Snoeren et al., 2010; Snoeren et al., 2011a), the higher than physiological dose of EB could in theory have triggered FLX-induced facilitation of sexual behavior. It was previously found that EB-priming of intact SERT knockout female rats mitigated reduced paracopulatory behavior shown in hormonally primed, ovariectomized SERT knockout females (Snoeren et al., 2010). Also when administered at adulthood, it has been shown that estradiol can attenuate the inhibiting effects of fluoxetine on female sexual behavior (Miryala et al., 2013). Another possible reason behind the differences in results, however, could also simply have been the timing of FLX exposure (perinatal versus postnatal) or the different strains that were used (Wistar versus Sprague-Dawley), as differences in strain sensitivity to FLX have been shown before (Maswood et al., 2008).

In our experiment, we used a seminatural environment in which rats are able to express all aspects of their natural behaviors. Mating patterns in groups of rats are quite different from the mating patterns observed in the traditional laboratory mating tests with pairs of rats or mice (Chu and Agmo, 2014, 2015b; Garey et al., 2002; McClintock, 1984; McClintock and Adler, 1978; McClintock and Anisko, 1982; McClintock et al., 1982; Snoeren et al., 2015). The use of the seminatural environment enabled us to evaluate the effects of perinatal SSRI exposure on natural mating behavior during the full behavioral estrus, in comparison to the short copulation tests previously used (Dos Santos et al., 2016; Rayen et al., 2014). Actually, we previously showed that in the same cohorts of Wistar rats, but with only a short observation period of 30 minutes, the FLX-females show slightly decreased levels of sexual activity (Houwing et al., 2019). We already hypothesized that the timing of observation could have influenced the findings, and that a proper evaluation of the full behavioral estrus should provide a clearer picture of the effects of perinatal FLX exposure on female sexual behavior. The current observations indeed show that the FLX-females (and CTR-females) that were not active during the previously scored 30-minute observation, copulated during the remainder of the behavioral estrus. Our findings are, therefore, the first to show the effects of perinatal SSRI exposure on the complete behavioral estrus. Not only did we not find any differences in behaviors over the full course of the behavioral estrus, also when the behavioral estrus was divided in different copulation bouts, FLX-females copulated in similar patterns as CTR-females within each of these bouts. No differences in female sexual behaviors were found in the *most active bout* or in for example the first or last copulation bout (data not shown). In addition, we found that there was no difference between CTR- and FLX-females in the onset or the duration of the behavioral estrus and *most active bout*. Therefore, we can be certain that the lack of effects was not caused by an effectively arbitrary timing of observations in which for example the FLX-females could have had a delay in start of copulation. Such differences would have been found in our experimental set-up.

The observation that both CTR- and FLX-females perform sexual behaviors in copulation bouts was rather peculiar. Our colleagues previously studied the full sexual behavioral patterns of male and female rats in a seminatural environment, with both intact and ovariectomized Wistar females (Chu and Agmo, 2014, 2015a, b; Le Moëne et al., 2020). While males copulated in bouts (Chu and Agmo, 2015b), female rats just showed one behavioral estrus per vaginal cycle (Chu and Agmo, 2014; Le Moëne et al., 2020). Their definition of a behavioral estrus was from the first lordosis to the last lordosis after which lordosis was absent for at least an hour. Although we started our observations with the same presumption, we quickly realized that all female rats start with a new behavioral bout at a later stage. Therefore, we defined *copulation bout* as the period from the first lordosis of a new bout to the last after which no lordosis was performed within an hour, and re-defined the behavioral estrus as the period from the first lordosis to the very last lordosis, after which no lordosis was seen until the end of the video recordings.

The reason behind this difference in sexual activity pattern remains unclear. Our hormonal priming resulted in the display of behavioral estrus of all females at the same time. One could argue that this might have caused a different behavioral pattern: the competition for mating could in theory have induced the display of new copulation bouts. (Chu and Agmo, 2014) indeed showed that when multiple females were in behavioral estrus at the same time, their estrus length increased and number of lordosis responses decreased in comparison to when only one female is sexually active. However, this did not result in the display of multiple copulation bouts. Another possibility would be that the others did not continue their observations after the one hour without lordosis, and more bouts could exist as well. Future research can hopefully explain the differences in sexual behavioral patterns found in our studies.

In terms of other social behaviors, no noteworthy differences were found in active social and conflict behaviors between CTR- and FLX-females. This finding is in contrast to another study that found increases in the time intact female rats spent on social investigation due to perinatal fluoxetine exposure (Gemmel et al., 2019). At the same, it also contrasts our own study in which we previously found slight decreases in the time spent on active social behavior in the seminatural environment, both in hormonally naïve and hormonally primed OVX female rats (Houwing et al., 2019). Moreover, we previously found that FLX-females have lower levels of conflict behavior compared to CTR-females, although conflicts happened rarely in general (Houwing et al., 2019). The most plausible explanation for these differences in finding is the fact that the females in the current experiment were observed during the behavioral estrus. Sexual interaction becomes the focus of the rats during this period, which could attenuate the effects of perinatal FLX on the other behaviors like social and conflict behavior.

For the current study the purpose for observing other social behaviors was to assure that potential differences in sexual behavior were not caused by abnormalities in active social or conflict behavior, as other have investigated the link between social and sexual behavior as well (Chu and Agmo, 2014, 2015b). Since no significant differences were found between CTR- and FLX-females, we can conclude that the lack of effects of perinatal FLX exposure on sexual behavior were not influenced by potential effects on active social or conflict behaviors.

At last, we also investigated the effects of perinatal FLX exposure on the expression pattern of ERα in the MPOM, VMNvl, BNST, MePD, and DRN. As mentioned before, the estrous cycle is under regulation of the HPG-axis, and there are indications that the serotonin system is involved in the development of this axis (Deneris and Gaspar, 2018; Millard et al., 2017). Since the activation and coordination of the HPG-axis is required for puberty establishment (Moran et al., 2013), it is then not surprising that repeated exposure to FLX *in utero* and during lactation was found to delay puberty onset in female rat offspring (Dos Santos et al., 2016). The effects on estrous cyclicity in adulthood, however, are unclear, with some studies finding regular patterns of estrous cyclicity (Dos Santos et al., 2016; Rayen et al., 2014) and another finding a tendency for elongated cycles in rats (Moore et al., 2015). Moreover, lesions of the DRN, the main source of serotonergic projections to the brain, in the prepubertal period induced a delay in puberty onset in female rats (Ayala et al., 1998). Overall, this suggests that the 5-HT exposure during early development can affect the onset of puberty and estrous cyclicity.

Serotonergic signaling from the DRN strongly innervates other parts of the brain (Phelix et al., 1992; Simerly et al., 1985), and these projections are known to also mediate hypothalamic estrogen receptor (ER) expression (Ito et al., 2014). The VMN, POA, MePD, and DRN are brain regions known for its role in the regulation in female sexual behavior (Arendash and Gorski, 1983; Hoshina et al., 1994; Kato and Sakuma, 2000; Kondo and Sakuma, 2005; Mathews and Edwards, 1977; Pfaff and Sakuma, 1979), and contain high levels of ERα (Laflamme et al., 1998; Yamada et al., 2009). Reduced expression levels of ERα in the VMN and POA have also shown to reduce paracopulatory and receptive behaviors in rats and mice (Musatov et al., 2006; Snoeren et al., 2015; Spiteri et al., 2010; Spiteri et al., 2012), while ERα in the BNST and MePD seems to play a minor role in female sexual behavior (Snoeren et al., 2015). If perinatal FLX exposure would affect the expression patterns of ERα in the brain, these regions would be the most logic targets. Our current data now showed that perinatal FLX exposure did *not* affect the number of ERα-IR cells in these regions. Although the serotonin system is involved in the development of the HPG axis and onset of puberty, our study suggests that increases in serotonin during early development does not have long-term consequences for the number of ERα-containing cells, at least not in the MPOM, VMNvl, BNSTpm, MePD, and DRN.

In addition, the lack of effect on ERα supports our behavioral findings that perinatal FLX exposure did not alter paracopulatory and receptive behaviors during their full behavioral estrus. It should be mentioned, though, that the ovariectomy itself could have affected the number of ERα-containing cells in the brain also in CTR-females. The literature is still unclear about the exact effect of ovariectomy on ERα expression patterns, but some studies have shown that ovariectomy can cause an increase in mRNA ERα levels in the uterus (Rosser et al., 1993), and ERα expression in the uterus and cerebral cortex (Mohamed and Abdel-Rahman, 2000). Others, however, have failed to detect an effect of ovariectomy on mRNA ERα levels in the arcuate nucleus and ventromedial hypothalamus (Simerly and Young, 1991). However, short- and long-term treatment with estrogens restored the effects found on ERα levels (Mohamed and Abdel-Rahman, 2000; Rosser et al., 1993). It therefore unlikely that our experimental procedure would have affected our findings, especially because both CTR- and FLX-females were ovariectomized. We have shown previously that differences in number of ERα-containing cells can be detected in ovariectomized rats (Snoeren et al., 2015), if perinatal FLX exposure would have altered the ERα expression in the brain regions of interest, we would most likely still have been able to detect a difference in number of ERα-containing cells.

## 5. Conclusion

Overall, we conclude that perinatal FLX exposure did not affect female rat sexual behavior in adulthood. CTR- and FLX-females had the same length and pattern of the full behavioral estrous period. No differences were found in paracopulatory and receptive behaviors during the full behavioral estrus, but also in the most active bout. Furthermore, no differences were found in the display of social and conflict behaviors, nor in the number of ERα-containing cells in several brain regions involved in sexual behavior. We conclude that although serotonin fluctuations during early development could affect neurodevelopmental effects, increases in serotonin levels during early development do not have long-term consequences for the regulation of female sexual behavior in adulthood.

## Acknowledgements

Financial support was received from Helse Nord #PFP1295-16, Norway. DJH was supported by the KNAW ter Meulen travel grant, the Netherlands. We also would like to thank Ragnhild Osnes, Carina Sørensen, Nina Løvhaug, Katrine Harjo, and Remi Osnes for their excellent care of the animals.

