## Supplemental materials for "Female rat sexual behavior is unaffected by perinatal fluoxetine exposure"

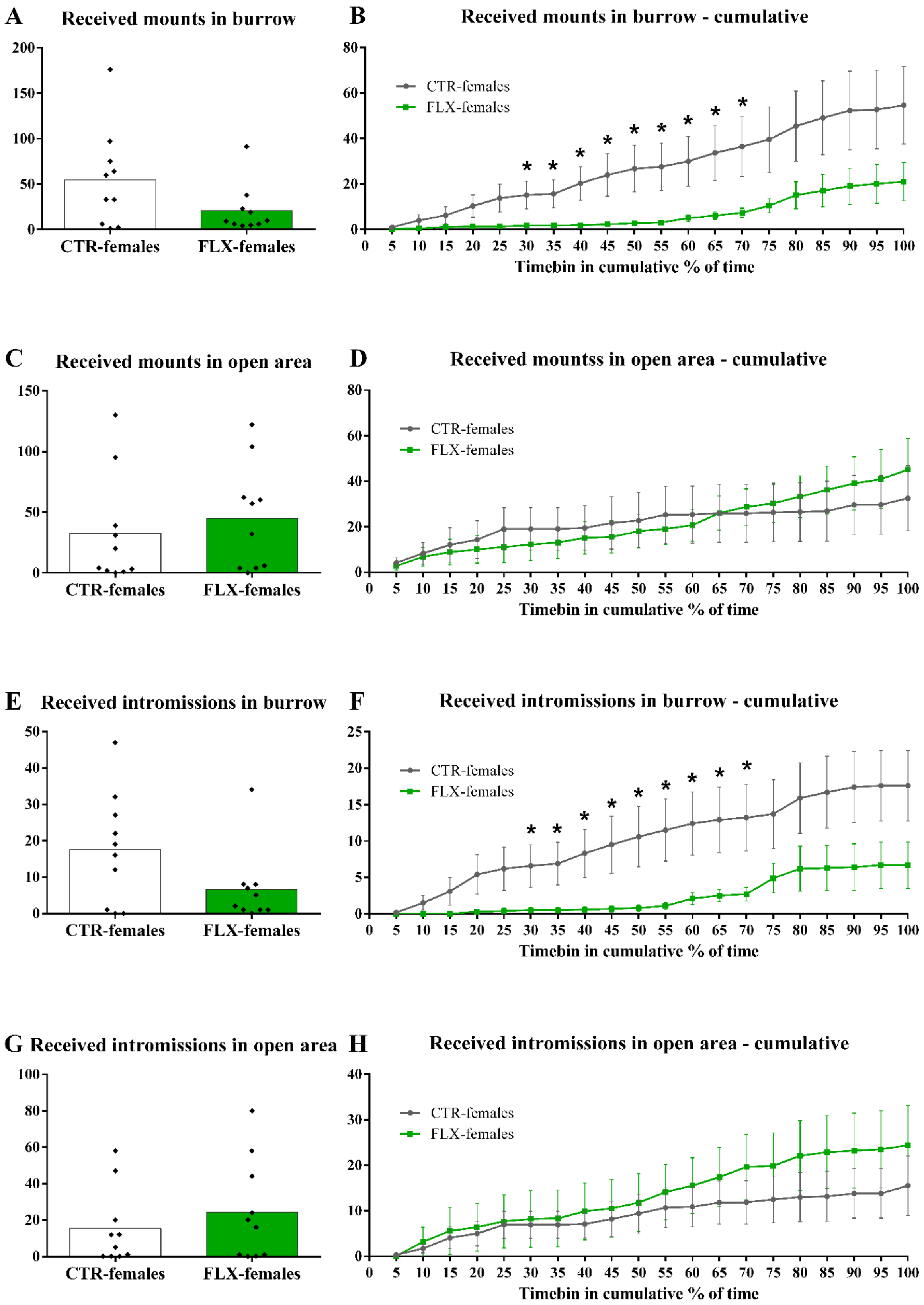
A
Received mounts in burrow
CTR-females
FLX-females
B
Received mounts in burrow - cumulative
CTR-females
FLX-females
Timebin in cumulative % of time
C
Received mounts in open area
CTR-females
FLX-females
D
Received mountss in open area - cumulative
CTR-females
FLX-females
Timebin in cumulative % of time
E
Received intromissions in burrow
CTR-females
FLX-females
F
Received intromissions in burrow - cumulative
CTR-females
FLX-females
Timebin in cumulative % of time
G
Received intromissions in open area
CTR-females
FLX-females
H
Received intromissions in open area - cumulative
CTR-females
FLX-females
Timebin in cumulative % of time

*Figure S1: Mounts and intromissions during the full behavioral estrus.*
*The data represents A) the number of received mounts in the burrow, B) the received mounts per cumulative 5% timebin of the behavioral estrus in the burrow, C) the number of received mounts in the open area, D) the received mounts per cumulative 5% timebin of the behavioral estrus in the open area, E) the number of received intromissions in the burrow, F) the received intromissions per cumulative 5% timebin of the behavioral estrus in the burrow, G) the number of received intromissions in the open area, H) the received intromissions per cumulative 5% timebin of the behavioral estrus in the open area. On the left, the data are shown with individual data points, with the bars representing the mean. On the right, the data are shown as mean±sem for CTR- and FLX-females. * $p<0.05$ compared to CTR-females.*

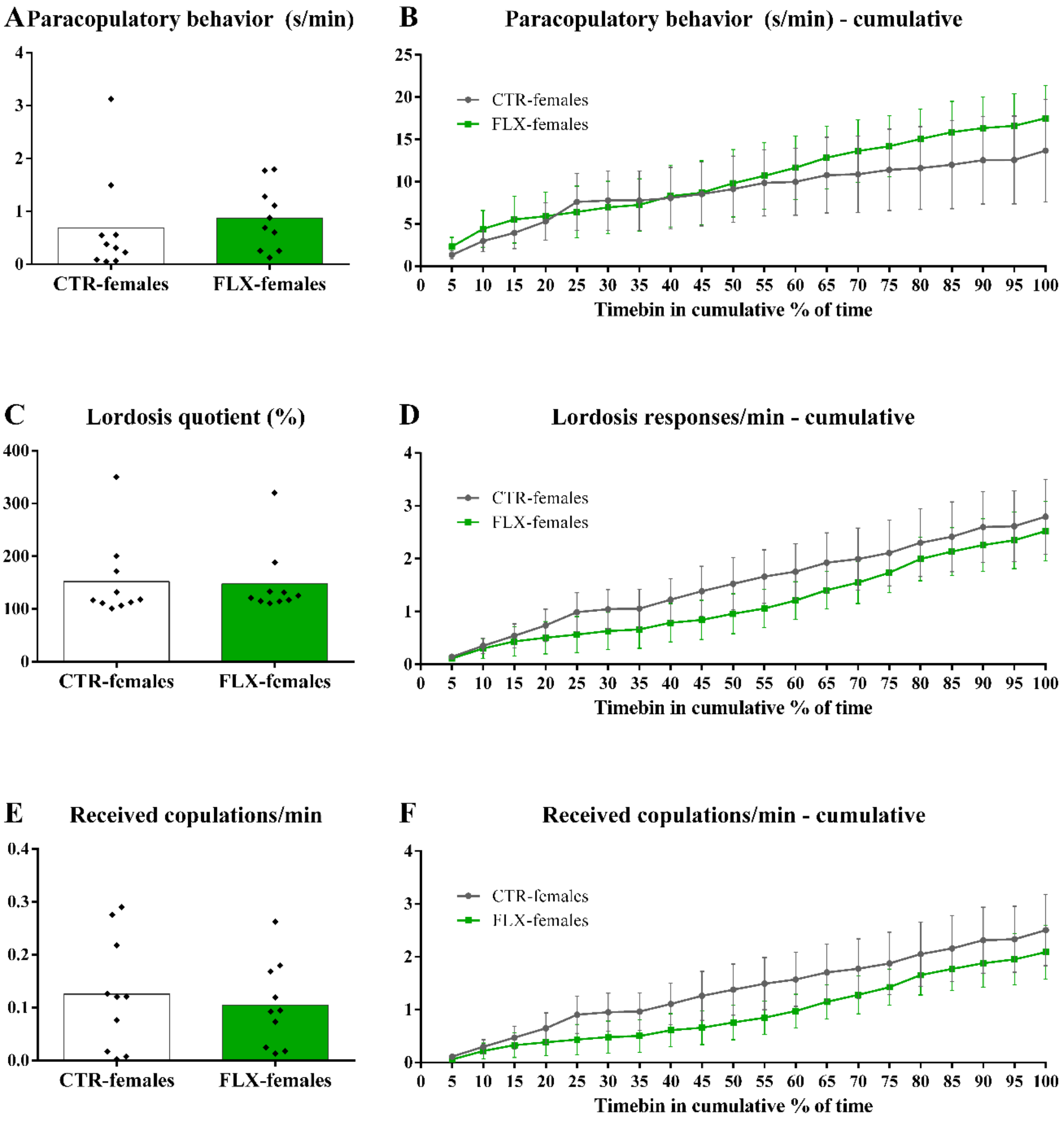


*Figure S2: Sexual behaviors per minute during the full behavioral estrus.*
*The data represents A) the time spent on paracopulatory behaviors in seconds per minute, B) the time spent on paracopulatory behaviors in seconds per minute per cumulative 5% timebin of the behavioral estrus, C) the lordosis quotient (lordosis responses/received copulations * 100%), D) the number of lordoses responses per minute per cumulative 5% timebin of the behavioral estrus, E) the number of received copulations (mounts+intromissions+ejaculations), and F) the number of received copulations per minute per cumulative 5% timebin of the behavioral estrus. On the left, the data are shown with individual data points, with the bars representing the mean. On the right, the data are shown as mean±sem for CTR- and FLX-females.*

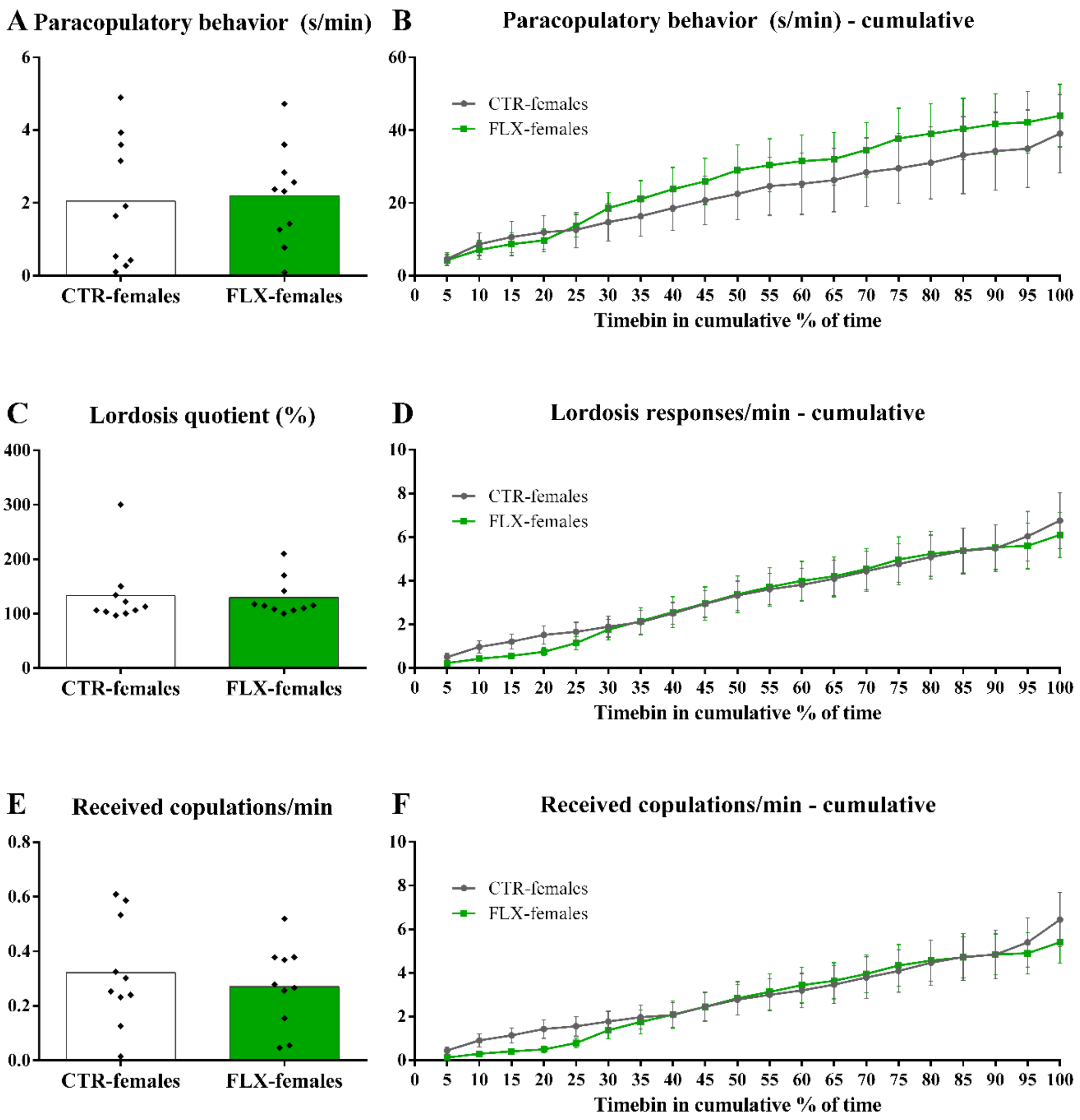


*Figure S3: Sexual behaviors per minute during the "most active bout".*
*The data represents A) the time spent on paracopulatory behaviors in seconds per minute, B) the time spent on paracopulatory behaviors in seconds per minute per cumulative 5% timebin of the "most active bout", C) the lordosis quotient (lordosis responses/received copulations * 100%), D) the number of lordoses responses per minute per cumulative 5% timebin of the "most active bout", E) the number of received copulations per minute (mounts+intromissions+ejaculations), and F) the number of received copulations per minute per cumulative 5% timebin of the "most active bout". On the left, the data are shown with individual data points, with the bars representing the mean. On the right, the data are shown as mean±sem for CTR- and FLX-females.*

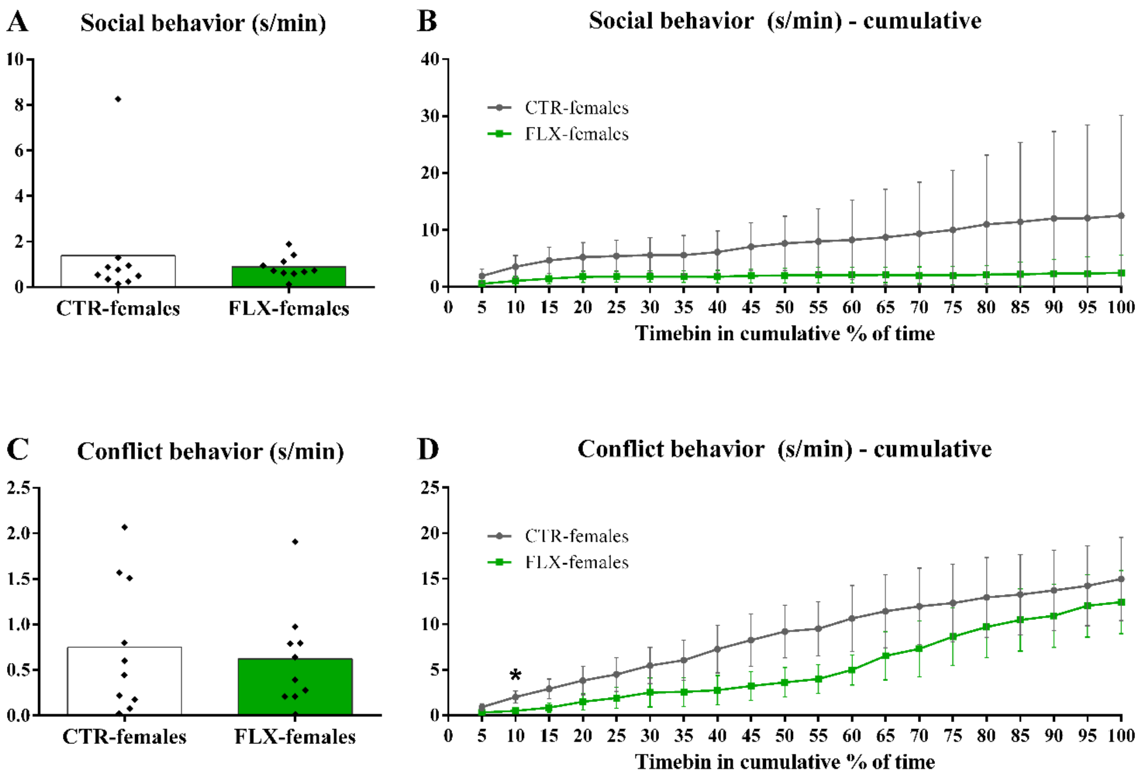


*Figure S4: Social behaviors per minute during the full behavioral estrus.*
*The data represents A) the time spent on social behaviors in seconds per minute, B) the time spent on social behaviors in seconds per minute per cumulative 5% timebin of the behavioral estrus, C) the time spent on conflict behaviors in seconds per minute, D) the time spent on conflict behaviors in seconds per minute per cumulative 5% timebin of the behavioral estrus. On the left, the data are shown with individual data points, with the bars representing the mean. On the right, the data are shown as mean±sem for CTR- and FLX-females. * p<0.05 compared to CTR-females.*

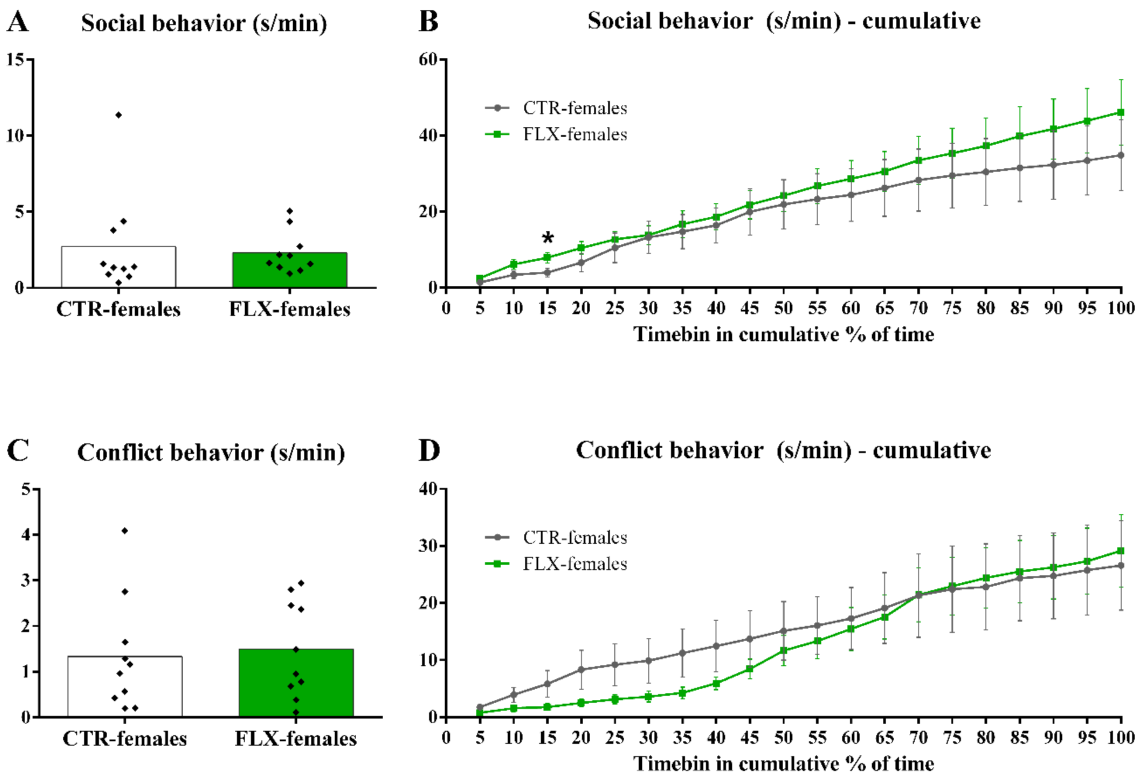


*Figure S5: Social behaviors per minute during the "most active bout".*
*The data represents A) the time spent on social behaviors in seconds per minute, B) the time spent on social behaviors in seconds per minute per cumulative 5% timebin of the "most active bout", C) the time spent on conflict behaviors in seconds per minute, D) the time spent on conflict behaviors in seconds per minute per cumulative 5% timebin of the "most active bout". On the left, the data are shown with individual data points, with the bars representing the mean. On the right, the data are shown as mean±sem for CTR- and FLX-females. * $p<0.05$ compared to CTR-females.*
