## Supplementary Table S1 for "Female rat sexual behavior is unaffected by perinatal fluoxetine exposure"

*The data represents the performed behaviors (number (#) or seconds (s)) in the total environment, in the burrow area, and in the open field.*

*Data are shown in mean±standard error of the mean during the behavioral estrus and during the most active bout.*

*No significant differences were found between CTR- and FLX females*

| | | Behavioral estrus | | | | | | Most active bout | | | | | |
|---|---|---|---|---|---|---|---|---|---|---|---|---|---|
| | | Total | | Burrow | | Open field | | Total | | Burrow | | Open field | |
| **Behavior** | **Rats** | **Mean** | **SEM** | **Mean** | **SEM** | **Mean** | **SEM** | **Mean** | **SEM** | **Mean** | **SEM** | **Mean** | **SEM** |
| Paracopulatory behavior (s) | CTR-females | 629.8 | 251.1 | 93.7 | 28.4 | 536.1 | 230.5 | 474.1 | 201.7 | 74.6 | 25.0 | 399.5 | 181.9 |
| | FLX-females | 799.0 | 170.3 | 89.9 | 33.1 | 709.1 | 183.8 | 545.2 | 121.3 | 69.2 | 28.8 | 476.1 | 131.2 |
| # Paracopulatory behaviors | CTR-females | 147.6 | 48.6 | 26.4 | 8.2 | 121.2 | 42.3 | 106.7 | 38.1 | 20.4 | 7.2 | 86.3 | 32.3 |
| | FLX-females | 200.2 | 41.7 | 23.0 | 7.4 | 177.2 | 44.4 | 136.9 | 29.7 | 18.1 | 6.9 | 118.8 | 31.7 |
| # Lordoses responses | CTR-females | 138.8 | 36.0 | 84.5 | 24.8 | 54.3 | 22.4 | 92.6 | 23.5 | 57.3 | 17.9 | 35.3 | 15.2 |
| | FLX-females | 122.4 | 28.4 | 38.0 | 13.1 | 84.4 | 26.2 | 82.1 | 16.9 | 25.9 | 12.9 | 56.2 | 16.6 |
| Lordosis quotient (%) | CTR-females | 151.8 | 24.2 | 161.2 | 30.3 | 132.7 | 23.5 | 133.3 | 19.3 | 107.8 | 5.8 | 166.7 | 36.1 |
| | FLX-females | 147.5 | 20.4 | 183.7 | 31.2 | 129.4 | 9.8 | 129.4 | 11.1 | 146.9 | 18.2 | 117.6 | 3.5 |
| # Total copulations (received) | CTR-females | 122.6 | 33.4 | 73.3 | 21.6 | 49.3 | 20.9 | 83.4 | 21.7 | 51.7 | 15.0 | 31.7 | 14.2 |
| | FLX-females | 100.2 | 25.1 | 28.2 | 12.0 | 72.0 | 22.6 | 70.2 | 15.5 | 21.2 | 12.0 | 49.0 | 14.6 |
| # Mounts (received) | CTR-females | 87.2 | 24.8 | 54.7 | 17.0 | 32.5 | 14.2 | 60.0 | 16.3 | 39.4 | 12.6 | 20.6 | 9.7 |
| | FLX-females | 66.2 | 16.1 | 21.1 | 8.5 | 45.1 | 13.8 | 42.8 | 8.9 | 15.3 | 8.4 | 27.5 | 7.6 |
| # Intromissions (received) | CTR-females | 33.1 | 9.2 | 17.6 | 4.8 | 15.5 | 6.6 | 21.8 | 6.2 | 11.8 | 2.9 | 10.0 | 4.6 |
| | FLX-females | 31.1 | 9.1 | 6.7 | 3.2 | 24.4 | 8.8 | 24.7 | 6.9 | 5.5 | 3.2 | 19.2 | 6.9 |
| # Ejaculations (received) | CTR-females | 2.3 | 0.9 | 1.0 | 0.4 | 1.3 | 0.7 | 1.6 | 0.8 | 0.5 | 0.2 | 1.1 | 0.6 |
| | FLX-females | 2.9 | 0.9 | 0.4 | 0.4 | 2.5 | 0.9 | 2.7 | 0.8 | 0.4 | 0.4 | 2.3 | 0.8 |
| Social behavior (s) | CTR-females | 1147.4 | 538.7 | 983.7 | 509.4 | 163.7 | 84.6 | 912.2 | 527.3 | 787.8 | 501.7 | 124.4 | 64.1 |
| | FLX-females | 817.2 | 149.3 | 668.2 | 150.4 | 149.1 | 31.1 | 588.5 | 140.5 | 498.9 | 143.5 | 89.7 | 24.6 |
| # Social behavior | CTR-females | 340.6 | 72.9 | 255.3 | 52.3 | 85.3 | 37.2 | 232.6 | 60.0 | 167.8 | 42.7 | 64.8 | 33.5 |
| | FLX-females | 383.9 | 51.2 | 261.3 | 42.8 | 122.6 | 25.2 | 264.3 | 40.7 | 190.0 | 36.0 | 74.3 | 20.3 |
| Sniffing others (s) | CTR-females | 1072.5 | 543.8 | 921.5 | 513.7 | 151.0 | 80.8 | 862.0 | 530.7 | 749.9 | 504.5 | 112.2 | 59.9 |
| | FLX-females | 707.1 | 130.8 | 567.8 | 132.3 | 139.3 | 30.5 | 512.7 | 127.1 | 428.8 | 129.9 | 83.9 | 23.9 |
| # Sniffing others | CTR-females | 318.9 | 71.4 | 238.3 | 50.6 | 80.6 | 35.6 | 216.4 | 58.5 | 156.1 | 41.6 | 60.3 | 31.7 |
| | FLX-females | 357.9 | 49.0 | 239.8 | 39.8 | 118.1 | 24.6 | 244.8 | 39.3 | 173.3 | 33.9 | 71.5 | 19.5 |
| Anogenitally sniffing (s) | CTR-females | 56.3 | 13.4 | 44.6 | 14.0 | 11.7 | 7.8 | 47.1 | 12.0 | 35.4 | 11.9 | 11.7 | 7.8 |
| | FLX-females | 78.9 | 27.0 | 70.1 | 24.8 | 8.8 | 3.1 | 62.8 | 27.3 | 57.4 | 25.0 | 5.3 | 2.8 |
| # Anogenitally sniffing | CTR-females | 19.5 | 4.1 | 15.2 | 4.0 | 4.3 | 2.5 | 15.2 | 3.7 | 11.0 | 3.3 | 4.2 | 2.5 |
| | FLX-females | 23.1 | 5.4 | 19.0 | 5.0 | 4.1 | 1.1 | 17.6 | 5.0 | 14.9 | 4.7 | 2.7 | 1.0 |
| Pursuing (s) | CTR-females | 0.0 | 0.0 | 0.0 | 0.0 | 0.0 | 0.0 | 0.0 | 0.0 | 0.0 | 0.0 | 0.0 | 0.0 |
| | FLX-females | 0.4 | 0.3 | 0.1 | 0.1 | 0.3 | 0.3 | 0.1 | 0.1 | 0.1 | 0.1 | 0.0 | 0.0 |
| # Pursuing | CTR-females | 0.0 | 0.0 | 0.0 | 0.0 | 0.0 | 0.0 | 0.0 | 0.0 | 0.0 | 0.0 | 0.0 | 0.0 |
| | FLX-females | 0.4 | 0.3 | 0.1 | 0.1 | 0.3 | 0.3 | 0.1 | 0.1 | 0.1 | 0.1 | 0.0 | 0.0 |
| Conflict behavior (s) | CTR-females | 684.0 | 195.8 | 677.3 | 196.5 | 6.7 | 2.6 | 379.9 | 131.5 | 373.9 | 131.9 | 6.0 | 2.7 |
| | FLX-females | 571.7 | 166.7 | 542.2 | 160.3 | 29.5 | 10.3 | 386.3 | 95.1 | 366.2 | 92.0 | 20.2 | 8.0 |
| # Conflict behavior | CTR-females | 186.5 | 52.6 | 182.0 | 53.3 | 4.5 | 1.9 | 98.2 | 25.1 | 94.4 | 25.8 | 3.8 | 2.0 |
| | FLX-females | 157.3 | 42.0 | 145.2 | 40.7 | 12.1 | 3.7 | 107.4 | 24.8 | 98.7 | 23.7 | 8.7 | 2.8 |
| Boxing/Wrestling (s) | CTR-females | 411.6 | 124.3 | 406.3 | 125.5 | 5.3 | 2.6 | 234.8 | 73.8 | 230.3 | 75.0 | 4.5 | 2.7 |
| | FLX-females | 333.3 | 78.7 | 305.6 | 73.0 | 27.7 | 10.3 | 235.6 | 70.8 | 217.2 | 69.0 | 18.4 | 8.2 |
| # Boxing/Wrestling | CTR-females | 125.2 | 37.9 | 121.2 | 38.6 | 4.0 | 1.9 | 66.6 | 16.2 | 63.3 | 17.0 | 3.3 | 2.0 |
| | FLX-females | 111.0 | 23.8 | 99.3 | 22.2 | 11.7 | 3.6 | 78.0 | 17.7 | 69.7 | 16.5 | 8.3 | 2.7 |
| Nose-off (s) | CTR-females | 271.4 | 103.6 | 270.9 | 103.5 | 0.5 | 0.5 | 144.1 | 66.3 | 143.5 | 66.3 | 0.5 | 0.5 |
| | FLX-females | 238.3 | 115.8 | 236.6 | 115.9 | 1.7 | 1.6 | 150.7 | 49.0 | 149.0 | 48.9 | 1.7 | 1.6 |
| # Nose-off | CTR-females | 60.9 | 19.8 | 60.7 | 19.8 | 0.2 | 0.2 | 31.2 | 10.3 | 31.0 | 10.2 | 0.2 | 0.2 |
| | FLX-females | 46.3 | 21.0 | 45.9 | 21.0 | 0.4 | 0.2 | 29.4 | 9.7 | 29.0 | 9.8 | 0.4 | 0.2 |
| Fighting (s) | CTR-females | 0.0 | 0.0 | 0.0 | 0.0 | 0.0 | 0.0 | 0.0 | 0.0 | 0.0 | 0.0 | 0.0 | 0.0 |
| | FLX-females | 0.0 | 0.0 | 0.0 | 0.0 | 0.0 | 0.0 | 0.0 | 0.0 | 0.0 | 0.0 | 0.0 | 0.0 |
| # Fighting | CTR-females | 0.0 | 0.0 | 0.0 | 0.0 | 0.0 | 0.0 | 0.0 | 0.0 | 0.0 | 0.0 | 0.0 | 0.0 |
| | FLX-females | 0.0 | 0.0 | 0.0 | 0.0 | 0.0 | 0.0 | 0.0 | 0.0 | 0.0 | 0.0 | 0.0 | 0.0 |
| Rejection (s) | CTR-females | 136.5 | 50.6 | 135.1 | 49.7 | 1.5 | 1.2 | 98.6 | 43.3 | 97.8 | 42.8 | 0.9 | 0.7 |
| | FLX-females | 153.4 | 44.4 | 145.6 | 43.2 | 7.9 | 4.1 | 95.0 | 38.5 | 91.9 | 38.9 | 3.2 | 1.8 |
| # Rejection | CTR-females | 39.7 | 11.9 | 38.5 | 11.9 | 1.2 | 1.0 | 25.2 | 7.6 | 24.6 | 7.6 | 0.6 | 0.5 |
| | FLX-females | 45.4 | 11.1 | 40.9 | 10.6 | 4.5 | 2.1 | 29.1 | 9.6 | 26.8 | 9.7 | 2.3 | 1.2 |
| Any other behavior (s) | CTR-females | 20148.5 | 4416.2 | 17486.7 | 4080.3 | 2661.7 | 640.0 | 13587.8 | 3094.9 | 11858.6 | 2969.3 | 1729.2 | 457.4 |
| | FLX-females | 19985.6 | 3021.4 | 13662.4 | 2071.1 | 6323.2 | 1625.1 | 12658.9 | 1368.6 | 8892.4 | 1370.1 | 3766.6 | 1058.6 |
| # Any other behavior | CTR-females | 721.1 | 145.2 | 528.9 | 115.7 | 192.2 | 60.3 | 460.9 | 90.6 | 330.6 | 70.7 | 130.3 | 47.1 |
| | FLX-females | 789.8 | 122.4 | 471.4 | 75.1 | 318.4 | 69.3 | 537.3 | 84.6 | 331.2 | 57.8 | 206.1 | 52.3 |
